# Emotion schema effects on associative memory differ across emotion categories at the behavioural, physiological and neural level

**DOI:** 10.1101/2022.03.04.482789

**Authors:** Monika Riegel, Marek Wypych, Małgorzata Wierzba, Michał Szczepanik, Katarzyna Jednoróg, Patrik Vuilleumier, Artur Marchewka

## Abstract

Previous behavioural and neuroimaging studies have consistently reported that our memory is enhanced for associations congruent or incongruent with the structure of our prior knowledge, termed as schemas. However, it remains unclear if similar effects exist if encoded associations are emotional. Do emotional schemas also facilitate learning and subsequent retrieval? Does it depend on the type of experienced emotions?

Using a novel face-word pair association paradigm combined with fMRI and eye-tracking techniques, we demonstrated and replicated in two independent studies that congruency with emotion schemas and emotion category interact to affect associative memory. Overall, emotion schemas facilitated memory for associative context, paralleled by the recruitment of left inferior frontal gyrus (IFG) during successful encoding of emotionally congruent vs. incongruent pairs. However, emotion schema effects differed across two negative emotion categories: disgust and fear, with disgust remembered better than fear. The IFG engagement was higher during successful encoding of congruent vs. incongruent pairs, but only in the case of disgust, suggestive of more semantic processing involved in learning disgust-related associations. On the contrary, the encoding of congruent vs. incongruent fear-related pairs was supported by activity in right fusiform gyrus (FG), suggesting greater sensory processing of faces. Successful memory formation for congruent disgust-related pairs was associated with a higher loading of pupil dilation component related to sympathetic activation, longer gaze time on words compared to faces, and more gaze switches between the two. This was reversed for fear-related pairs where the faces attracted more attention, as reflected by longer gaze time (compared to words).

Overall, our results at the behavioural, physiological, and neural level converge to suggest that emotional congruency influences memory similar to semantic schemas. However, encoding processes and neural effects vary depending on emotion category, reflecting the differential role of semantic processing and visual attention processes in the modulation of memory by disgust and fear.

## Introduction

In everyday life, we constantly form new memories of complex events. For instance, when having a conversation with a good friend, we memorize multiple, associated aspects of this conversation and its context - not only the semantic meaning of the words we use, but also the facial expressions of our friend, as well as emotions we both convey. How do we form and retrieve these associations with emotional context? Does it depend on the type of emotions we experience? And does this mechanism differ when learned associations are consistent or contradictory to our previous experience of such conversations?

Memory is often studied as if the new information was imposed on a tabula rasa, but considerable research evidence suggests otherwise. An effective memory system requires that we integrate new information with prior knowledge (Bartlett, 1932; Bein et al., 2015a; DeWitt, Knight, Hicks, & Ball, 2012; JW Alba, 1983; Kole & Healy, 2007). Such higher-level knowledge structures, termed as “schemas”, reflect commonalities among experiences and serve as general reference templates to which new information can be compared (Amer, Giovanello, Nichol, Hasher, & Grady, 2019; Bein, Reggev, & Maril, 2020). Indeed, a large body of research confirms that prior knowledge, or schemas, influence learning of new associations (DeWitt et al., 2012; Kole & Healy, 2007; Reder, 2013) in at least two different ways (Greve, Cooper, Tibon, & Henson, 2019; Quent, Greve, & Henson, 2021). First, information *congruent* with schemas can be better integrated into our knowledge structures. It was demonstrated that we are better at remembering associations between words, objects and scenes if they are semantically congruent (Bein et al., 2015b; van Kesteren et al., 2013). This effect is supported by the inferior frontal gyrus (IFG) and paralleled with a rapid shift from initial processing of memories in the medial temporal lobe (MTL) towards long-term memory representations in the medial prefrontal cortex (mPFC) (Brod, Lindenberger, Werkle-Bergner, & Shing, 2015; Preston & Eichenbaum, 2013a; Tse et al., 2007). On the other hand, a memory advantage and higher attentional engagement (Mudrik, Deouell, & Lamy, 2011) also occur for novel associations *incongruent* with schemas (van Kesteren, Ruiter, Fernández, & Henson, 2012). This effect is supported by the hippocampus (HC) and other MTL regions, possibly due to greater saliency and intensified processing of prediction errors (Chang & Sanfey, 2009; Greve, Cooper, Kaula, Anderson, & Henson, 2017). This way, we preferentially remember person descriptions that violate appearance-based first impressions and trustworthy faces paired with negative behavior descriptions (Bell, Mieth, & Buchner, 2015; Cassidy & Gutchess, 2015). However, it remains unclear what can constitute a “schema”.

One important candidate factor, also well known to modulate memory of associations (Cahill & McGaugh, 1998; Hamann, 2001; Kensinger, Garoff-Eaton, & Schacter, 2006), is emotion. Appraisal theories (Moors, 2020) have proposed that emotions are canonical responses to situations ancestrally linked to survival and constructed from multiple perceptual, mnemonic, and conceptual ingredients (Russell, 2003). These particular responses should be driven by specific patterns of organism-environment interactions called “emotion schemas” (Izard, 2007). Recent neuroimaging studies demonstrate that emotion schemas are embedded both in sensory input (Kragel, Reddan, LaBar, & Wager, 2019) and mPFC representations (Philipp Paulus, Ian Charest, & Roland Benoit, n.d.). Moreover, emotion expression and recognition was shown to depend on acquired conceptual knowledge (Bertoux et al., 2020). Specific emotion schemas should differ across emotion categories, such as disgust and fear. For example, an image of someone approaching a threatening animal evokes rapid responses related to fear, whereas an image of someone eating contaminated food evokes disgust responses. Both fear and disgust are negative, arousing, and motivate avoidance (Krusemark & Li, 2011; Woody, 2000), and their primary function is to protect us from threat or contamination. To this end, one needs to form durable memories of events and contexts evoking these essential emotions – that is, specific emotion schemas. Indeed, there is evidence showing that disgust and fear can differentially influence memory at the behavioral level (Chapman, Johannes, Poppenk, Moscovitch, & Anderson, 2012; Croucher, Calder, Ramponi, Barnard, & Murphy, 2011; van Hooff, Devue, Vieweg, & Theeuwes, 2013; van Hooff, van Buuringen, El M’rabet, de Gier, & van Zalingen, 2014), possibly mediated by distinct patterns of visual attention (Fink-Lamotte, Svensson, Schmitz, & Exner, 2021a; Schienle, Potthoff, Schönthaler, & Schlintl, 2021) and physiological responses (Critchley et al., 2005; De Jong, van Overveld, & Peters, 2011; Susskind et al., 2008; Woody, 2000). Our recent study showed distinct engagement of key brain regions supporting emotional memory, i.e. amygdala (AMY) and MTL areas during encoding and reinstatement of disgust and fear (Riegel et al., 2022). However, it remains unknown how memory modulation by disgust and fear is related to mechanisms typical to schemas such as congruency and incongruency effects, as well as what are the concomitant physiological and neural mechanisms.

One possibility is that emotions indeed provide a specific type of schema or a particular form of saliency for learned associations. We approach this possibility disregarded by previous research by referring to everyday communication. There is clearly a higher probability of co-occurrence of verbal utterances and facial expressions related to the same emotion, which in turn activates specific reactions and expectations. If this co-occurence could constitute an emotion schema, at the behavioral level we should observe typical mnemonic benefits for newly learned, emotionally congruent associations. A difficult binding of emotionally incongruent words and faces should be reflected by an increased number of gaze fixations (Frank, Montaldi, Wittmann, & Talmi, 2018). It could also imply distinct neural mechanisms, whereby the reliance on prior knowledge (in the case of congruent associations) should be supported by left inferior frontal gyrus (IFG) and medial prefrontal cortex (mPFC). On the other hand, medial temporal lobe (MTL) regions such as the hippocampus (HC), as well as sensory cortex such as right fusiform gyrus (FG) should be engaged in forming more complex relational binding for information violating prior knowledge (in the case of incongruent associations) (Brod et al., 2015; van Kesteren et al., 2013).

Another possibility is that emotional characteristics of newly learned associations modulate memory with distinct attentional, physiological and neural mechanisms, overriding typical schema-related effects. That is, whether verbal utterances and facial expressions are related to the same or different emotions (i.e. fitting or not our emotion schemas), we would not observe any congruency or incongruency effects on memory. Instead, we could expect to see more automatic emotion effects, driven, for example, by a direct activation of the noradrenergic system due to arousal (Clewett & Murty, 2019; Sterpenich et al., 2006). A recent electrophysiological study using a dot probe task with images presented as distractors showed that threatening images, but not semantically incongruent scenes and objects, capture attention automatically and rapidly (Furtak et al., 2020). Likewise, we would expect to see the emotion-specific effects during encoding at the level of visual attention - how often and for how long we look at words vs, faces, for different predictions see: (Fink-Lamotte, Svensson, Schmitz, & Exner, 2021b; Schienle et al., 2021; Schienle, Übel, Gremsl, Schöngassner, & Körner, 2016); and physiological reactions - pupil response reflecting arousal (Aston-Jones & Cohen, 2005b), pupil temporal dynamics reflecting sympathetic vs. parasympathetic activation (Clewett, Gasser, & Davachi, 2020a; Johansson, Pärnamets, Bjernestedt, & Johansson, 2018a). Moreover, we would expect to observe different neural mechanism of encoding for disgust- vs. fear-related pairs (distinct engagement of AMY and MTL regions (Riegel et al., 2022).

Finally, it is possible that both existing emotion schemas, and emotional characteristics of newly learned information influence memory, and their effects are not exclusive. Integrating new emotional information into existing emotion schemas can differ between disgust and fear. For instance, when seeing a friend’s facial expression of disgust, its congruency with the verbal communication could affect our memory more than in the case of fear, although no existing literature suggests a specific direction to this hypothesis. Labeling facial expressions related to disgust and fear, inherent to our experimental task, was shown to enhance encoding of contextual information (Barrett & Kensinger, 2010), yet differences between them were not explored. Our recent findings showed that these two emotions could imply different sensory and semantic processing of events and their context (Riegel et al., 2022), suggesting that they could differentially modulate congruency and incongruency effects. Therefore, memory encoding might disclose not only emotion-specific effects of the event content, but also additive or interactive effects of an emotional congruency with the context, both described above.

To investigate these issues, we run two independent experiments with the same experimental paradigm combined with a multi-method approach necessary to test our hypotheses. We deployed eye-tracking (ET) and pupillometry (Exp. 1) to investigate the engagement of visual attention and physiological processes, and functional magnetic resonance (fMRI; Exp. 2) to study the underlying brain mechanisms. In both experiments we asked human participants to learn associations between words (as messages) and faces (as interlocutors); each conveying either disgust, fear, or being emotionally neutral; arranged in emotionally congruent or incongruent conditions. For instance, the word *knife* (fear-related) was paired with a face expressing fear (congruent) or disgust (incongruent). During retrieval, participants were cued with previously presented (or new) words alone and asked to retrieve the emotion expressed by the associated face. By analyzing gaze direction during encoding (fixations and switches between words and faces), we assessed how associations were initially formed. Further, by analyzing pupil dilation changes, we could elucidate the role of sympathetic and parasympathetic activation during encoding. Finally, using fMRI with a univariate activation analysis, we investigated the role of schema and emotion processing during encoding of these word-face associations. We found associative (word-face) memory facilitation due to emotional congruency, which affected memory similar to typical effects of prior knowledge and schemas. However, this effect varied depending on emotion category, which reflects differential semantic and visual attention processing implicated in memory modulation by disgust and fear. Together, we showed at the attentional, physiological, and neural levels that when forming memories of emotional context, we rely both on emotional schemas and emotional content of learnt associations.

## Materials and methods

### Participants

Native Polish speakers with no history of any neurological illness or treatment with psychoactive drugs, right-handed and with normal or corrected-to-normal vision, took part in both experiments. They were mostly students and young professionals living in Warsaw, with at least secondary education. In Exp. 1, the sample of n = 31 participants included only females (aged 22-29, M = 25.3, SD = 3.1). In Exp. 2, the sample of n = 37 participants included only females as well (aged 22-29, M = 25.3, SD = 3.1). Four subjects were excluded from the fMRI analyses due to excessive head motion (> 7 mm with a voxel size 3.5 mm). Only 18 subjects had >= 10 trials per experimental condition per run and only those were included in the main fMRI analyses (for an extended fMRI analysis see: results and supplementary materials, Fig. S8). All participants provided informed consent and were financially compensated. The local Research Ethics Committee at Faculty of Psychology, University of Warsaw approved the experimental protocol of the study.

### Stimuli

We selected 420 words from the Nencki Affective Word List (NAWL) (Riegel et al., 2015). Based on the available ratings of valence, arousal, as well as basic emotions (Wierzba et al., 2015), we selected 140 words eliciting disgust, 140 words eliciting fear, and 140 neutral words. Emotional stimuli were controlled for other affective scales (valence, arousal, and intensity of other basic emotions). Eighty out of 140 words in each emotion category (disgust, fear, neutral) were used in both encoding and retrieval phases as targets (old), whereas the remaining 60 words were additionally included as lures (new) in the retrieval phase.

During encoding, these words were paired with 240 (60 per run) faces selected from standardized datasets of affective faces (FACES (Ebner, Riediger, & Lindenberger, 2010), n = 144; KDEF (Lundqvist, Flykt, & Ohman, 1998), n = 66; and WSEFEP(Olszanowski et al., 2014), n = 30): 80 (20 per run) faces expressing disgust, 80 (20 per run) faces expressing fear, and 80 (20 per run) neutral faces. The faces were selected from multiple datasets to ensure the selection of unique faces expressing specific emotion categories, counterbalanced in terms of sex, age and emotion across datasets.

The words were pseudorandomly assigned as either targets (240) or lures (180), counterbalanced in terms of emotion category. Target words and faces were pseudorandomly assigned to experimental conditions, as either: emotionally congruent: 40 (10 per run) disgust word + disgust face; 40 (10 per run) fear word + fear face, 80 (20 per run) neutral word + neutral face; or emotionally incongruent: 40 (10 per run) disgust word + fear face, 40 (10 per run) fear face + disgust word). Words and faces were randomly paired within these conditions. Note that neutral pairs were always congruent and served as a basic control condition.

The order of experimental trials in both encoding and retrieval phases was pseudorandom under the following constraints: no more than 3 consecutive trials of the same emotion category (disgust, fear, neutral), no more than 3 of the same part of speech (noun, verb, adjective), no more than 4 old (targets) or new (lures) in the retrieval phase. The final set of Polish words used as stimuli included the following examples (here: translated to English) for disgust: *bug, rotten, drainpipe, rat, shit, stinking*; for fear: *suffocate, knife, bomber, lurk, faint, collision*; and for neutral: *circle, sentence, brush, magnet, liquid, stone*. Supplementary materials, Table S1, provides a full list of word-face pairs categorized according to experimental conditions, together with the presentation order during each run.

### Study design and experimental paradigm

In both experiments (ET, Exp. 1 and fMRI, Exp. 2) we used the same experimental paradigm, consisting of four experimental sessions of encoding (6.5 min) and immediate retrieval (11-16 min), followed by rest breaks (Fig. 1a). Before entering the scanner, subjects were given the details of the scanning procedure and performed a brief practice session (comprising 10 word - face pairs) either in front of the computer (Exp. 1) or in a mock MRI scanner (Exp. 2). The experimental procedure was programmed using the Presentation software (Neurobehavioral Systems, Inc., Albany, CA, USA). In Exp. 2, the procedure was displayed on an MR-compatible high-resolution LCD monitor positioned at the back of the scanner. Subjects observed the stimuli through a mirror system placed on a MR coil.

**Fig. 1.**
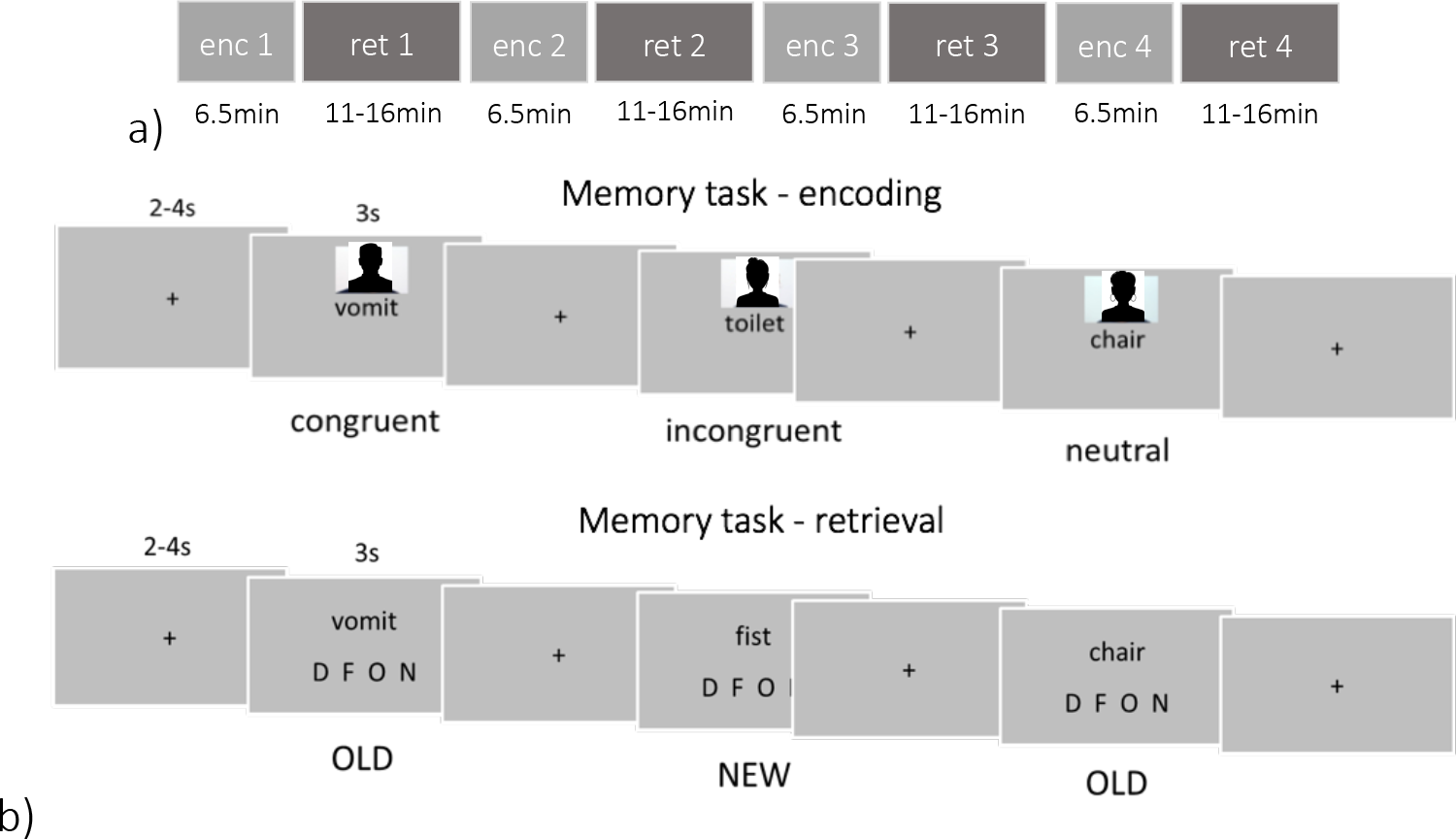
a) Overview of experimental procedure with four sessions of encoding and immediate retrieval; b) Example trial structure for all the emotion and congruence conditions during encoding and retrieval; enc – encoding, ret – retrieval, D - disgust, F - fear, O - other (neutral), N – new In each experimental session, immediately after encoding, the subjects performed a retrieval phase (Fig. 1b). During a cued associative retrieval task, participants were presented with 60 words from the encoding list (targets) and 45 new words (lures), that is 105 trials in total (3s + jittered fixation cross for 2-4s), and asked to determine (3s) what was the emotional expression of the face previously associated with each word. They could also report that they had not seen the word before, that is it was new. Additionally, for 20% of correctly retrieved old facial emotion, participants were presented with a face and asked to indicate whether this face was previously associated with a given word or not (2.5s). This task was added to ensure the subjects stayed motivated and focused on the task. The instruction was formulated as follows: “You will be presented with words expressing different emotions or neutral words. Your task will be to indicate, if you can remember having seen them, what was the emotion of the previously associated face [D - disgust, F - fear, O - other (neutral)] or indicate that the word is new [N - new]”.

Each of four experimental sessions started with encoding, as depicted in Fig. 1b. The participants were presented (3s + jittered fixation cross for 2-4s) with 60 word-face pairs (20 disgust-related congruent and incongruent, 20 fear-related congruent and incongruent and 20 neutral congruent). They were instructed to imagine the word as a message expressed by a person whose face was presented (to create a communicative context), and memorize as many of these pairs as possible. The instruction was formulated as follows: “Try to memorize as many pairs of words and faces as possible. In order to do so, imagine each word as a message coming from the accompanying face.”

### Behavioral data analysis

#### Memory performance

First, memory for emotional and neutral stimuli was assessed as performance in the cued associative retrieval task. Of note, the chance level in this task was 25%, given 4 possible responses. Specifically, the retrieval rate was calculated as a proportion of correctly retrieved facial emotions previously associated with an old word. To analyze memory as a function of facial emotion category (3 levels: disgust, fear, neutral) and congruency with emotion schemas (2 levels: congruent, incongruent) as within-subject factors, a repeated-measures analysis of variance (rm ANOVA) was performed on the retrieval rate (proportion of correct responses) data.

#### Reaction times

Reaction times (RT) in the memory task performance were also analyzed to investigate the speed of memory decisions and memory accessibility (Lifanov, Linde-Domingo, & Wimber, n.d.; Sheldon, Williams, Harrington, & Otto, 2020) due to emotion and congruency with emotion schemas. Previous research indicates that RT can be used to assess and quantify the confidence in memory tasks (Weidemann & Kahana, 2016). We performed rm ANOVA with emotion (3 levels: disgust, fear, neutral), congruency (2 levels: congruent, incongruent), and correctness (2 levels: correct, incorrect) as within-subject factors.

The mean number and proportion of trials with standard deviations for all experimental conditions used in the analyses of Exp. 1 (with ET) and Exp. 2 (with fMRI) is presented in Table S2. All the reported ANOVAs were performed with the Greenhouse-Geisser adjustments to the degrees of freedom when the assumption of sphericity was violated. Post hoc comparisons were performed using the Bonferroni adjustment for multiple comparisons. The results were visualized with the use of Python-based Seaborn package (Waskom, 2021) and Statannot package (https://github.com/webermarcolivier/statannot).

### Eye-tracking and pupillometry data acquisition and preprocessing

The study was performed using the Tobii Pro TX300 infra-red eye tracker. We used a 23- inch TFT monitor with a resolution of 1920 × 1080 pixels to display the stimuli while gaze behavior was simultaneously recorded. The system tracked both eyes to a rated accuracy of 0.5° with a sampling rate of 300 Hz. The participants were tested individually, facing the eye-tracker monitor at a distance of approximately 60 cm. The experimenter sat next to the participant to control the computer without interfering with viewing behavior.

A calibration procedure consisting of 9 points displayed on a plain, grey background was performed before each of four sets of encoding and retrieval runs. If needed, the procedure was repeated until the calibration result was considered valid (no more than 0.5° accuracy).

Participants’ eye movements were recorded both during encoding and retrieval phases, but only the former will be presented in the current manuscript focused on encoding mechanisms. The gaze direction data was preprocessed using custom code implemented in MATLAB R2018b (Mathworks, Inc., Natnick, MA, USA) and the ILAB software (Gitelman, 2002). The regions of interest (ROI) were defined as two non-overlapping rectangles centered on the presented face and word, each with a size 20% larger than the average size of stimuli. Fixations were detected by the dispersion-threshold identification (I-DT) algorithm (Salvucci & Goldberg, 2000), under threshold of minimal duration of 100ms and a size of 0.5 degree of visual angle. Continuous data was split into meaningful segments, that is, trial-locked and epoched from 0 until 3000 ms according to each trial duration.

The pupil data was preprocessed following guidelines available in literature(Kret & Sjak- Shie, 2019; Mathôt, Fabius, Van Heusden, & Van der Stigchel, 2018; Zénon, 2017) with custom code implemented in MATLAB R2018b (Mathworks, Inc., Natnick, MA, USA). First, the artifacts such as: eye blinks, signal dropouts (pupil > than .02mm) and extreme variation (pupil change > than .38 mm within a 20-ms intervals (Snowden et al., 2016), were removed. Then, the data was smoothed using S-G coefficients, and the missing data was linearly interpolated. Subjects with < 50% valid pupil data in all runs of encoding or retrieval (n = 3) were removed from the final analysis (final sample for pupil analyses: n = 28). The averaged pupil diameter was adjusted (detrended) independently of the trial type to account for linear trends possibly present due to overall subjects’ fatigue. Then, we extracted the trial-locked data, epoched according to trial duration from 0 until 3000 ms. We used the average pupil size during 100 ms preceding each trial onset(Karatekin, Couperus, & Marcus, 2004) (fixation cross, gray screen) as baseline. To calculate baseline-corrected pupillary responses, we subtracted the baseline pupil size from the average pupil size at stimulus individually (trial-by-trial) and then averaged across trials. As a result, for each participant and each experimental condition, we had 900 samples (300 Hz x 3 seconds of trial length) of baseline-corrected pupil dilation data.

### Eye-tracking and pupillometry data analysis

#### Eye movements analysis

Gaze direction data was used to assess visual attention and its impact on mnemonic processes. The statistical analyses were conducted on the average of the following measures: percent (%) of gaze time in each ROI, number of fixations in each ROI, number of gaze switches between ROIs. To analyse gaze direction as a function of emotion of facial expressions (3 levels: disgust, fear, neutral), and congruency with emotion schemas (2 levels: congruent, incongruent) and ROI (word, face) as within-subject factors, a repeated-measures analysis of variance (rm ANOVA) was performed on gaze data (gaze time, number of fixations and number of switches). All the reported ANOVAs were performed with the Greenhouse-Geisser adjustments to the degrees of freedom when the assumption of sphericity was violated. Post hoc comparisons were performed using the Bonferroni adjustment for multiple comparisons. The results were visualized with the use of Python-based Raincloud plots package (Allen et al., 2021) and Statannot package (https://github.com/webermarcolivier/statannot).

#### Pupil dilation - time windows analysis

Pupil dilation data was used to assess physiological arousal and cognitive effort (D Kahneman, 1966; Joshi & Gold, 2020; Joshi, Li, Kalwani, & Gold, 2016), as well as its relationship with memory (Kucewicz et al., 2018). The first set of statistical analyses was conducted on the average baseline-corrected pupil dilation change (relative to the baseline, that is after subtracting 100 ms preceding each trial) during pre-defined time windows. Based on previous memory and emotion research, these time windows of interest were equal to trial duration - 0 –3 s (Johansson, Pärnamets, Bjernestedt, & Johansson, 2018b). To analyze how arousal or cognitive effort was modulated by emotion of facial expressions (3 levels: disgust, fear, neutral), congruency with emotion schemas (2 levels: congruent, incongruent) and correctness (2 levels: correct, incorrect) as within-subject factors, a repeated-measures analysis of variance (rm ANOVA) was performed on the baseline-corrected pupillary changes. The results were visualized with the use of Python-based Raincloud plots package (Allen et al., 2021), Statannot package (https://github.com/webermarcolivier/statannot) and custom Matlab scripts.

#### Pupil dilation – temporal PCA

To further understand how emotion and congruency modulated pupil dilation, and how these effects could reflect ongoing physiological reactions related to sympathetic and parasympathetic activation, we performed a temporal principal component analysis (PCA) on the preprocessed pupil data, adapted from previous work (Clewett et al., 2020a). First of all, average baseline-normed pupil dilation was computed for each experimental condition (emotion, congruency, correctness). We averaged these values across participants to reduce the amount of input variables (300 variables; 30 participants x 10 conditions) and to control for noisy, spontaneous pupil changes on individual trials. We included all 3s trial-locked pupil samples (900 samples per condition in total) as dependent measures in the PCA.

First, we performed a visual inspection based on a scree plot (depicting eigenvalues of factors) and selected reliable components. Five components with an eigenvalue > 10 in total explained more than 97% of variance. A Varimax rotation was performed on the components output by the temporal PCA. An unrestricted PCA using the covariance matrix with Kaiser normalization and Varimax rotation was used to generate maximal loadings on each component with minimal overlap with other components. The resulting loadings reflect the correlated, temporally dynamic patterns of pupil dilation elicited by stimuli presentation during encoding. Factor loadings with eigenvalues > 1 were analyzed in subsequent analyses (Kaiser criterion).

Because the PCA was data-driven and agnostic to condition, we were able to then examine the relative contributions of loading patterns for each pupil component according to the different emotion and congruency conditions. To analyze temporal characteristics of pupil dilation as a function of emotion of facial expressions (3 levels: disgust, fear, neutral), congruency with emotion schemas (2 levels: congruent, incongruent) and correctness (2 levels: correct, incorrect) as within-subject factors, repeated-measures analysis of variance (rm ANOVA)

was performed on the component loadings (comp1, comp2, comp3, comp4, comp5). As this analysis was exploratory, no adjustments were made for multiple comparisons (Clewett et al., 2020a). The results were visualized with the use of Python-based Seaborn package (Waskom, 2021) and Statannot package (https://github.com/webermarcolivier/statannot).

#### Relationship between gaze direction, pupil dilation and memory across conditions

Next, we aimed test whether the observed modulation of visual attention and pupil responses by emotions and congruency could be related to memory success. Thus, we performed an exploratory correlation analysis (Pearson’s r correlation coefficients) between the gaze direction measures, pupil dilation change and pupil component loadings per condition with retrieval rate per condition. In order to directly compare the resulting correlation coefficients across emotion categories, we used r-to-z transformation and comparison for independent samples (Lenhard & Lenhard, 2014). As these analyses were exploratory, no adjustments were made for multiple comparisons.

Visual complexity control

Finally, in order to control for the possible effects of visual characteristics of images (faces) used in the experiment, we performed a metadata analysis using a custom Python script. We calculated the following measures: luminance (an average pixel intensity of the gray-scaled image), contrast (SD of all luminance values, see: (Bex & Makous, 2002)) and entropy (a statistical measure of the randomness/ noisiness of an image). These measures for each face image were compared between emotions (3 levels: disgust, fear, neutral) with one-way ANOVA to indicate if it was equal for the experimental conditions (see: Table S5 in supplementary materials).

### fMRI data acquisition and preprocessing

Magnetic resonance data was acquired on a 3T MAGNETOM Trio TIM system (Siemens Medical Solutions) equipped with a whole-head 32-channel coil. The following images were acquired within a single subject scanning: structural localizer image, structural T1-weighted (T1w) image (TR: 2530 s, TE: 3.32 ms, flip angle: 7°, PAT factor = 2, voxel size 1 x 1 x 1 mm, field of view 256 mm, volumes: 1, TA = 6:03min), field map (TR: 423 ms, TE1: 5:19 ms, TE2: 7.65 ms, flip angle: 60°, voxel size 3.5 x 3.5 x 3.5 mm, field of view 224 mm, volumes: 1, TA: 0:56 min), 8 series of functional EPI images (TR: 2500 ms, TE: 27 ms, flip angle: 90°, PAT mode = none, voxel size 3.5 x 3.5 x 3.5 mm, field of view 224 mm, volumes: 153 in encoding runs, and 270-383 in retrieval runs, TA: 6:29 min encoding runs and 16:04 min retrieval runs), functional localizer (TR: 2500 ms, TE: 27 ms, flip angle: 90°, PAT mode = none, voxel size 3.5 x 3.5 x 3.5 mm, field of view 224 mm, volumes: 37, TA: 1:39 min). The whole scanning of encoding and retrieval phases took approximately 120 minutes, independent of participants’ speed of responding (as trials had fixed duration), but varied due to memory performance and the presentation of additional face questions (on 20% of correct responses). Only fMRI data from encoding will be reported in the current manuscript.

At the initial step of MRI data preprocessing, the DICOM series were converted to NIfTI using the MRIConvert (ver. 2.0.7; Lewis Center for Neuroimaging, University of Oregon). Then, imaging data was preprocessed using Statistical Parametric Mapping (SPM12; Wellcome Department of Cognitive Neuroscience, University College London, London, UK) running under Matlab 2013b (Mathworks, Inc., Natnick, MA, USA). Preprocessing started with the correction of functional images for distortions related to magnetic field inhomogeneity and correction for motion by realignment to the first acquired image. Then, structural (T1w) images from single subjects were co-registered to the mean functional image and segmented into separated tissue classes (grey matter, white matter, cerebrospinal fluid) using the default tissue probability maps (TPM). Structural and functional images were normalized to the MNI space and resliced to preserve the original resolution. Finally, functional images were smoothed with the 6 mm FWHM Gaussian kernel.

### fMRI data analysis

#### Univariate activation analysis – whole brain

A standard univariate activations analysis approach was used to examine the difference at each voxel between the average activity across experimental conditions (emotion, congruency, correctness). MRI data was recorded both during encoding and retrieval phases, but only encoding results will be reported here. fMRI data was initially analyzed with SPM12 based on general linear models (GLMs). At the subject level, each single event was modelled with onset and duration corresponding to the presentation of a word-face pair, together with a response given afterwards (as it was not sufficiently separated in time from a pair presentation). To account for movement-related variance, six nuisance regressors were included as representing the differential of the movement parameters from the realignment. A default high-pass filter cutoff of 128 seconds was used to remove low-frequency signal drifts. Data was convolved with a standard canonical hemodynamic response function (HRF) to approximate the expected blood-oxygen-level dependent (BOLD) signal.

At the subject level, the interaction between emotion, congruency, and memory performance was examined for all the encoding and retrieval runs in one single model. Functional volumes from all the encoding and retrieval runs were split across conditions along the factors of emotion of facial expressions *(DIS - disgust, FEA - fear, NEU - neutral)*, congruency with emotion schemas *(con - congruent, incon - incongruent)* and correctness *(corr - correct, incorr - incorrect)*. Experimental conditions representing brain activity during encoding were the following: DIS con corr, DIS con incorr, DIS incon corr, DIS incon incorr, FEA con corr, FEA con incorr, FEA incon corr, FEA incon incorr, NEU con corr, NEU con incorr. Experimental conditions representing brain activity during retrieval were the following: DIS con corr, DIS con incorr, DIS incon corr, DIS incon incorr, FEA con corr, FEA con incorr, FEA incon corr, FEA incon incorr, NEU con corr, NEU con incorr, NEW corr, NEW incorr. Altogether 4 runs x 10 encoding conditions and 4 runs x 12 retrieval conditions were included in the model. The number of correct trials per experimental condition and per run depended on individual memory performance and varied between subjects (ranging from 10 to 35). Only subjects having at least 10 correct trials in each experimental condition in each run were further included in the group-level analysis (n = 18). However, we made attempts to increase the sample size and performed another analysis, this time modeling only the best runs (>= 10 trials per condition) from 33 subjects (for the results see: supplementary materials, Table S4 and Fig. S8).

At the group level, ANOVAs were performed by including within-subject factors of emotion and congruency with schemas, as well as a subject factor. The flexible factorial design (Ashburner et al., 2014) was used because of the abovementioned variability in the number of trials among subjects and possible subject effects. Activation maps were created for the interaction effects of emotion of facial expressions (DIS, FEA) and congruency with emotion schemas (con, incon) with an exclusive masking procedure allowing us to identify selective increases produced by successful but not unsuccessful encoding (Gauvin, De Baene, Brass, & Hartsuiker, 2016; Hofstetter, Achaibou, & Vuilleumier, 2012; B. P. Staresina, Duncan, & Davachi, 2011) (determined by subsequent incorrect retrieval). The analysis of interaction effects was based on (Henson, 2015), with the contrasts were created as on p. 479. Note that all possible combinations of conditions lead to two overall contrasts, presented in Fig. 7. The neutral condition was always congruent, therefore not included in this analysis.

If not stated otherwise, in all analyses, we applied a voxel-wise height threshold of p < .1 (uncorrected) combined with a cluster-level extent threshold of p < .05, corrected for multiple comparisons using the Monte Carlo simulation (3dClustSim algorithm) for the whole brain analyses (nn = 2, two-sided, p = .001, alpha = .05), in line with the most recent AFNI recommendations to reduce the false positive rate (Cox, Chen, Glen, Reynolds, & Taylor, 2017). A small volume correction (SVC) (Worsley et al., 1996) was applied using anatomical masks corresponding to specific regions-of-interest (ROI), selected according to a priori hypotheses about their engagement in emotion and memory interactions, listed below. Within each of these ROIs, we considered activations whose effects survived the SVC correction at the voxel level. This type of correction for multiple comparisons was previously used in studies on emotion and memory due to the small volume of subcortical structures of interest (Domínguez-borràs et al., 2017; Richter, Cooper, Bays, & Simons, 2016). The coordinates of significant effects were reported in the Montreal Neurological Institute (MNI) space and labeled according to Automated Anatomical Labeling (AAL2) (Rolls, Joliot, & Tzourio-Mazoyer, 2015a) atlas with the use of bspmview (http://www.bobspunt.com/bspmview). Results were visualized with the use of Python-based Nilearn plotting function (https://nilearn.github.io/plotting/index.html) and MRIcroGL (http://www.mccauslandcenter.sc.edu/mricrogl/home).

#### Region-of-Interest localization and analyses

Based on the available literature, several regions of interest (ROI) were selected to further test specific hypotheses concerning differences in how emotions of facial expressions and congruency with emotion schemas affects successful encoding (followed by correct retrieval). The ROI analysis was performed using the MarsBaR toolbox (Brett, Anton, Valabregue, & Poline, 2002). Contrast estimate values were extracted based on the subject-level SPM models used in the first univariate ANOVA analysis mentioned above for the following regions: bilateral amygdala (AMY), bilateral hippocampus (HC), left inferior frontal gyrus (IFG), and right fusiform gyrus (FG). Although mPFC was also extensively studied with relation to schema and congruency effects, it was not consistently demonstrating differences in univariate activation during encoding between congruent and incongreunt items (Bein, Reggev, & Maril, 2014b; Brod & Shing, 2018; van Kesteren et al., 2013). Thus, we did not have clear hypotheses and did not include mPFC in the primary ROI analyses. However, it was added as an exploratory analysis and added to supplementary materials, Fig. S6. Anatomical masks for the ROI analysis were specified according to the AAL2 (Rolls et al., 2015a) atlas implemented in the WFU PickAtlas (Maldjian, Laurienti, Kraft, & Burdette, 2003) toolbox (version 3.0.5).

To examine how the engagement of these ROIs during encoding is modulated by our experimental conditions, we first performed a rm ANOVA on the contrast estimates extracted from each ROI, with emotion of facial expressions (3 levels: disgust, fear, neutral), congruency with emotion schemas (congruent, incongruent) and correctness (2 levels: correct, incorrect) as within-subject factors. To unpack these effects, we run a post hoc analysis of simple effects restricted to correctly encoded pairs, with two emotion categories (disgust, fear) and schema congruency (congruent, incongruent). We used paired t-tests for dependent samples to directly compare the mean contrast estimates between different levels of emotion and congruency factors for all ROIs.

All the reported ANOVAs were performed with the Greenhouse-Geisser adjustment to the degrees of freedom when the assumption of sphericity was violated. Post hoc comparisons were performed using the Bonferroni adjustment for multiple comparisons. The results were visualized with the use of Python-based Seaborn package (Waskom, 2021) and Statannot package (https://github.com/webermarcolivier/statannot).

## Results

### Emotion and congruency with schemas modulate memory performance

In Exp. 1 (with ET), memory of facial emotions in response to word cues was tested immediately after encoding. Correct facial emotion memory differed depending on both emotion expression of the face (disgust, fear) and congruency with the word (congruent, incongruent), with a significant interaction effect of these two factors [F(1,30) = 61.415, p < .001, η2 = .672], Fig 2A. Of note, in each experimental condition, the average retrieval rate was higher than the chance level (that is - 25% with 4 possible responses in the cued associative retrieval task). Faces from pairs congruent with emotion schemas were generally better retrieved (p < .001). Among congruent pairs, disgust faces were retrieved more often than fear (p < .001), whereas among incongruent pairs fear faces were retrieved more often than disgust (p = .011). This result pattern could also mean that disgust words (present in: congruent disgust and incongruent fear conditions) served as a more effective retrieval cue than fear words, possibly enhancing associations with both a disgusted (congruent; p < .001) and a fearful (incongruent; p = .021) face expression. However, these differences were also driven by an interaction effect [F(1,30) = 4.548, p = .041, η2 = .132], meaning both emotion and congruency modulated memory.

**Fig. 2.**
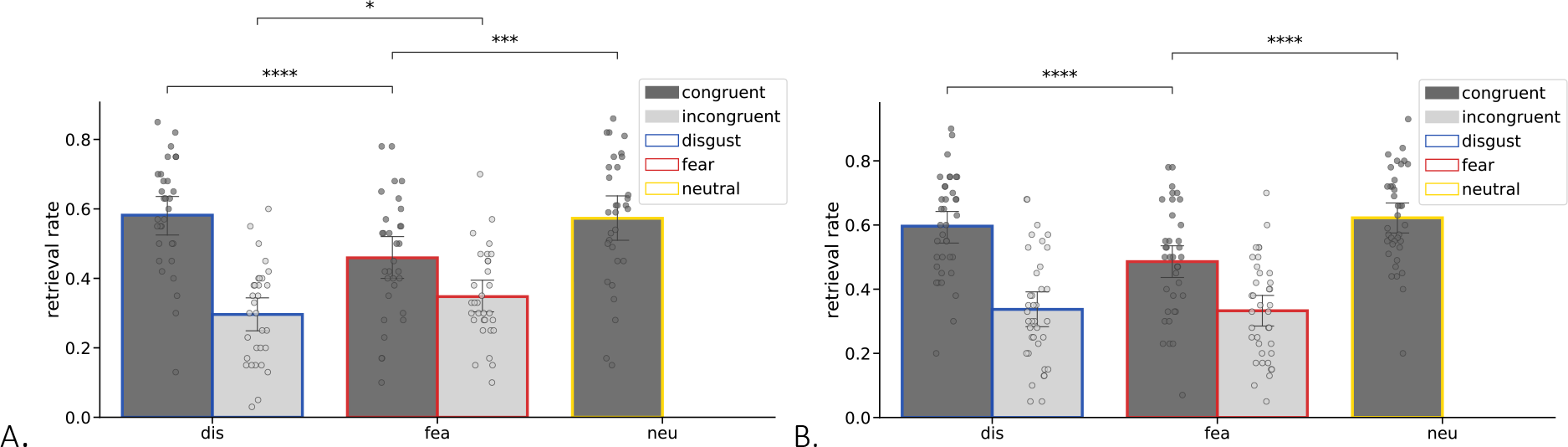
Retrieval rate (proportion of correctly retrieved facial emotions previously associated with old words) across emotion and congruency conditions in A. Exp. 1, and B. Exp. 2. *Frame colors and labels on horizontal (x) axis indicate the emotion of associated faces to be retrieved: blue = disgust, red = fear, yellow = neutral; bar colors indicate the congruency with emotion schemas: dark grey – congruent, light grey – incongruent; error bars represent one standard deviation, * p < .05, *** p < .001, **** p < .0001*.

In comparison, the neutral faces (always emotionally congruent pairs) were retrieved better than congruent fear faces (p = .001) but equally to congruent disgust faces (p = .336), with a significant effect of emotion across the three old congruent conditions [F(2,30) = 13.948, p < .001, η2 = .317], as shown in Fig. 2A. This result indicates a mnemonic cost due to fear, both compared to disgust, and to neutral baseline, a baseline that was easier to remember being always congruent. Correct rejection of new words (i.e. lures during retrieval) did not differ between emotion categories of word cues [F(2,30) = 1.926, p = .160, η2 = .051], Fig. S1A.

The results described above were largely replicated in a separate cohort in Exp. 2 (with fMRI). Exp. 2 again revealed an interaction effect between emotion type (disgust, fear) and congruency with schemas (congruent, incongruent) [F(1,36) = 18.074, p < .001, η2 = .334]) on correct retrieval. Like in Exp. 1, congruent disgust faces were retrieved more often than congruent fear faces (p < .001), while there was no significant difference between emotions for incongruent pairs, Fig. 2B. Also similar to Exp. 1, disgust words presented during retrieval were a better retrieval cue than fear words, but only in congruent pairs (“congruent disgust” condition; p < .001) but not incongruent pairs (“incongruent fear” condition; p = .868), with an interaction effect [F(1,36) = 9.743, p = .004, η2 = .213]. Again, like in Exp. 1, fear faces were retrieved less often than neutral (p < .001) and disgust faces (p < .001), with no difference between neutral and disgust (p = 1.000), and a significant effect of emotion across all three congruent conditions, F(2,72) = 19.786, p < .001, η2 = .355, Fig. 2B. Note that like in Exp. 1, the neutral condition in Exp. 2 was always congruent. Correct rejection of new words did not differ between emotions related to word cues [F(2,64) = 2.69, p < .079, η2 = .078], Fig. S1B.

Given that in the fMRI analyses of data from Exp. 2 (with fMRI) we could only include subjects having at least 10 trials per experimental condition (n = 18), we repeated the behavioural analyses for this subset of participants. Likewise, we also analysed a subset subjects having >= 10 trials per experimental condition in Exp. 1 (with ET; n = 16). The results of these additional analyses are mostly consistent with the results of the whole sample analyses presented above, and are included in supplementary materials and Fig. S4.

### Emotion and congruency with schemas modulate reaction time during retrieval

In Exp. 1, reaction times (RT) during the retrieval task differed depending on emotion of facial expressions, congruency with emotion schemas, and response correctness (three-way interaction effect [F(2,60) = 11.125, p = .002, η2 = .271]), Fig. 3A and Fig. S2A. To unpack this interaction, we ran a post hoc analysis of simple effects that showed differences for pairs congruent with emotion schemas only. Correct retrieval of congruent disgust pairs was faster than congruent fear (p = .002), whereas incorrect retrieval (the proportion of incorrectly retrieved facial emotions previously associated with old words) of congruent disgust pairs was slower than incorrect congruent fear (p = .002). There were no effects of correctness and emotion for incongruent pairs.

**Fig. 3.**
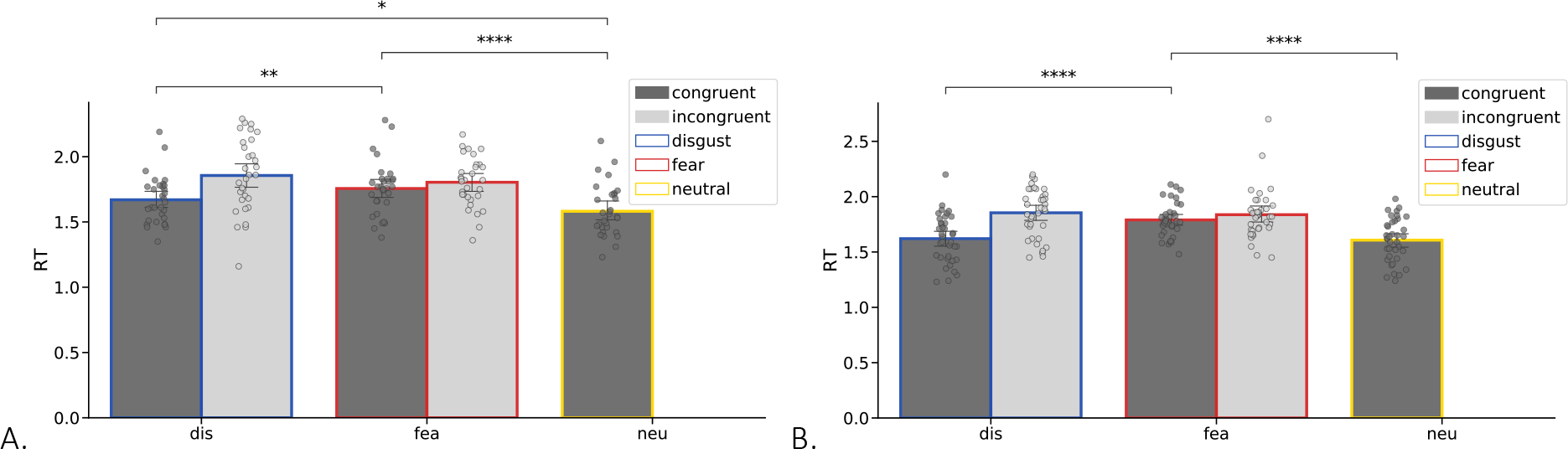
Reaction times (RT) in the retrieval task for correct retrieval of facial emotions, measured in seconds from the word cue onset until response, across the different emotion and congruency conditions in A. Exp. 1, and B. Exp. 2. *Frame colors and labels on horizontal (x) axis indicate the emotion of associated faces to be retrieved: blue = disgust, red = fear, yellow = neutral; bar colors indicate the congruency with emotion schemas: dark grey – congruent, light grey – incongruent; error bars represent one standard deviation; * p < .05, *** p < .001, **** p < .0001. Note that (RT) in the retrieval task for incorrect retrieval of facial emotions was presented in Fig. S3A and B*.

Likewise, when considering only old pairs congruent with emotion schemas (Fig. 3A and Fig. S2A), RTs for retrieval differed depending on emotion category and correctness (interaction effect: [F(2,60) = 10.278, p < .001, η2 = .255]), with correct retrieval of fear faces being significantly slower than both disgust (p < .006) and neutral faces (p < .001), and correct retrieval of disgust faster than neutral faces (p = .032). Correct rejection of new words was also modulated by emotion [F(2,64) = 5.613, p = .006, η2 = .149], faster for neutral than disgust-related (p = .01) and, to a lesser degree, fear-related words (p = .057), Fig. S3A. Overall, the RT results specific for correct retrieval mirrors the memory performance results.

Replicating results from Exp. 1 (Fig. 3B and Fig. S2B) RTs in Exp. 2 differed depending on emotion of retrieved facial expressions, congruency with emotion schemas, and correctness (three-way interaction effect [F(1,36) = 8.156, p = .007, η2 = .185]), Fig. 3B and Fig. S2B. Like in Exp. 1, post hoc analysis showed that correct retrieval of congruent disgust faces was faster than congruent fear (p < .001), whereas incorrect retrieval of incongruent disgust was slower than incongruent fear (p = .022). Replicating Exp. 1, for congruent (old) pairs alone, correct RTs showed an interaction effect of emotion and correctness [F(2,72) = 12.822, p < .001, η2 = .263], with correct retrieval of fear being significantly slower than both disgust (p < .001) and neutral faces (p < .001), and incorrect retrieval of neutral faces being faster than both disgust (p < .001) and fear (p < .001). Again, like in Exp.1, RT of correct rejection of new words differed between emotions [F(2,70) = 5.716, p = .007, η2 = .140], faster for neutral than disgust-related (p = .004) and fear-related words (p = .001), Fig. S3B. Overall, the RT results in Exp. 2 replicated the RT results from Exp. 1, and likewise echoed the memory performance results.

### Visual attention to stimulus pairs depends on emotion and schema congruency

In Exp. 1, gaze direction was recorded to assess how visual attention differences during encoding was modulated by emotion of facial expressions and congruency with emotion schemas. We first focused on the percentage of gaze time spent by participants on each ROI in the display (word and face). We found its modulation by facial emotions and congruency with emotion schemas both in ROI1 (face) (interaction effect [F(1,28) = 10.521, p = .003, η2 = .273]) and ROI2 (word) (interaction effect [F(1,28) = 16.852, p < .001, η2 = .376]), Fig. 4A and B. Participants looked longer at ROI1 (face) for congruent fear pairs than both congruent disgust (p < .001) and incongruent fear pairs (p = .004). There was no difference between incongruent pairs. Conversely, they looked shorter at ROI2 (word) for congruent fear than congruent disgust (p = .036) and incongruent fear pairs (i.e., when the word cue was related to disgust; p < .001). Globally, across conditions people also spent more time looking at faces than words, as expected given the task instructions to further retrieve facial expressions (p < .001), driven by an interaction with ROI when added to the analysis [F(1,30) = 17.565, p < .001, η2 = .369].

**Fig. 4.**
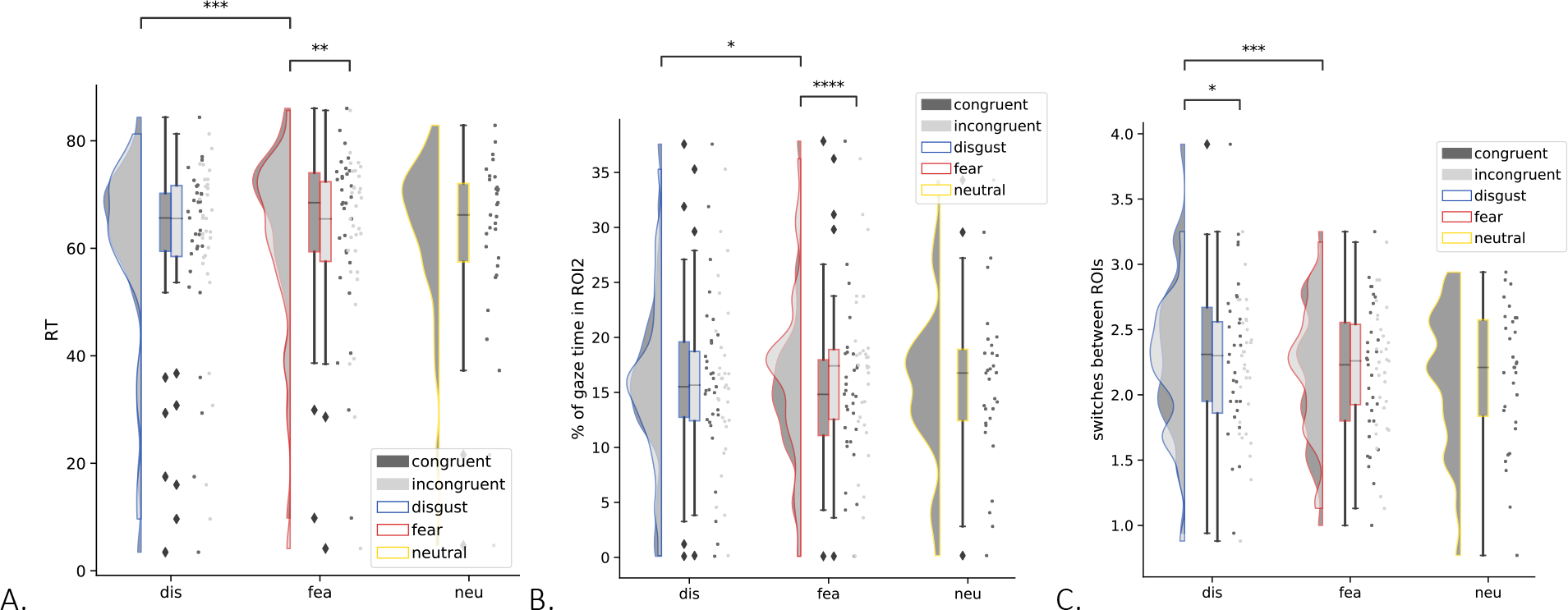
Gaze direction during encoding reflecting visual attention distribution as a function of facial emotion and congruency with emotion schemas. A. Percentage of overall gaze time spent in ROI1 (face); B. Percentage of overall gaze time spent in ROI2 (word); C. Number of gaze switches between ROIs. *Frame colors and labels on horizontal (x) axis indicate the emotion of associated faces to be retrieved: blue = disgust, red = fear, yellow = neutral; bar colors indicate the congruency with emotion schemas: dark grey – congruent, light grey – incongruent. Each dot represents one participant, the probability density of the data is shown as raincloudplots. Boxplots indicate the median and interquartile range (IQR, i.e., distance between the first and third quartiles). The lower and upper hinges correspond to the first and third quartiles (the 25th and 75th percentiles). The upper whisker extends from the hinge to the largest value no further than 1.5* IQR from the hinge. The lower whisker extends from the hinge to the smallest value at most 1.5* IQR of the hinge. Error bars indicate ±1 SEM; * p < .05, ** p < .01, *** p < .001*.

Another measure of interest was the number of fixations on each ROI. We observed a similar pattern of results, with emotion of facial expressions and congruency with emotion schemas modulating the number of fixations in both ROI1 (face; interaction effect [F(1,28) = 4.068, p = .053, η2 = .127]) and ROI2 (word; interaction effect [F(1,28) = 13.579, p = .001, η2 = .327]), Fig. 4C. Similar to the previous analysis, we found more fixations on the face when participants encoded congruent fear than congruent disgust pairs (p = .006), and no difference for incongruent pairs. On the other hand, we observed more fixations on words when encoding congruent disgust (p = .003) and incongruent fear pairs (with a disgust-related word; p = .022) compared with congruent fear pairs.

Finally, we also examined the number of saccades between ROIs (gaze switches), assumed to reflect the process of comparing different stimuli or integrating pieces of information (for instance, learning associations) (Eckstein, Guerra-Carrillo, Miller Singley, & Bunge, 2017). Gaze switches between ROIs were computed as a function of emotion of facial expressions and congruency with emotion schemas, and again showed an interaction effect [F(1,28) = 7.772, p = .009, η2 = .217]. Participants switched gaze more often between faces and words when encoding congruent disgust compared to fear pairs (p <. 001) and compared to incongruent disgust pairs (p = .044), but there was no significant difference between incongruent disgust and fear pairs. Fig 4C.

### Facial emotions and emotion schemas modulate pupil dynamics during encoding

Pupil diameter is controlled by brain systems involved in autonomic functions (Eckstein et al., 2017) and may reflect the engagement of physiological arousal and cognitive effort, which modulate attention and memory (Clewett, Huang, Velasco, Lee, & Mather, 2018; Sterpenich et al., 2006). Therefore, in Exp. 1, we analyzed the mean baseline-corrected pupil size in time-windows covering the trial length (0-3s) and compared changes on successfully encoded trials as a function of emotion, congruency and correctness (Fig. 5A). We found that both facial emotions and schema congruency influenced pupil size (interaction effect [F(1,27) = 12.057, p = .002, η2 = .309], driven by greater dilation when encoding incongruent than congruent fear pairs (p < .001) (i.e., faces paired with a disgust- rather than fear-related word), and also greater dilation during encoding of congruent disgust than congruent fear pairs (p = .005). There was no interaction with correctness [F(1,27) = 1.350, p = .255, η2 = .048]. Next, in a separate analysis we verified if the observed effect could be driven by the presence of disgust-related words (as in incongruent fear and congruent disgust pairs), and we found the main effect of emotion [F(1,27) = 12.057, p = .002, η2 = .309], and a trend of the main effect of congruency [F(1,27) = 3.486, p = .073, η2 = .114]. However, these modulatory effects overlapped with a global light-driven constriction evoked by the visual display onset and were clearly changing over time (Fig. 5B).

**Fig. 5.**
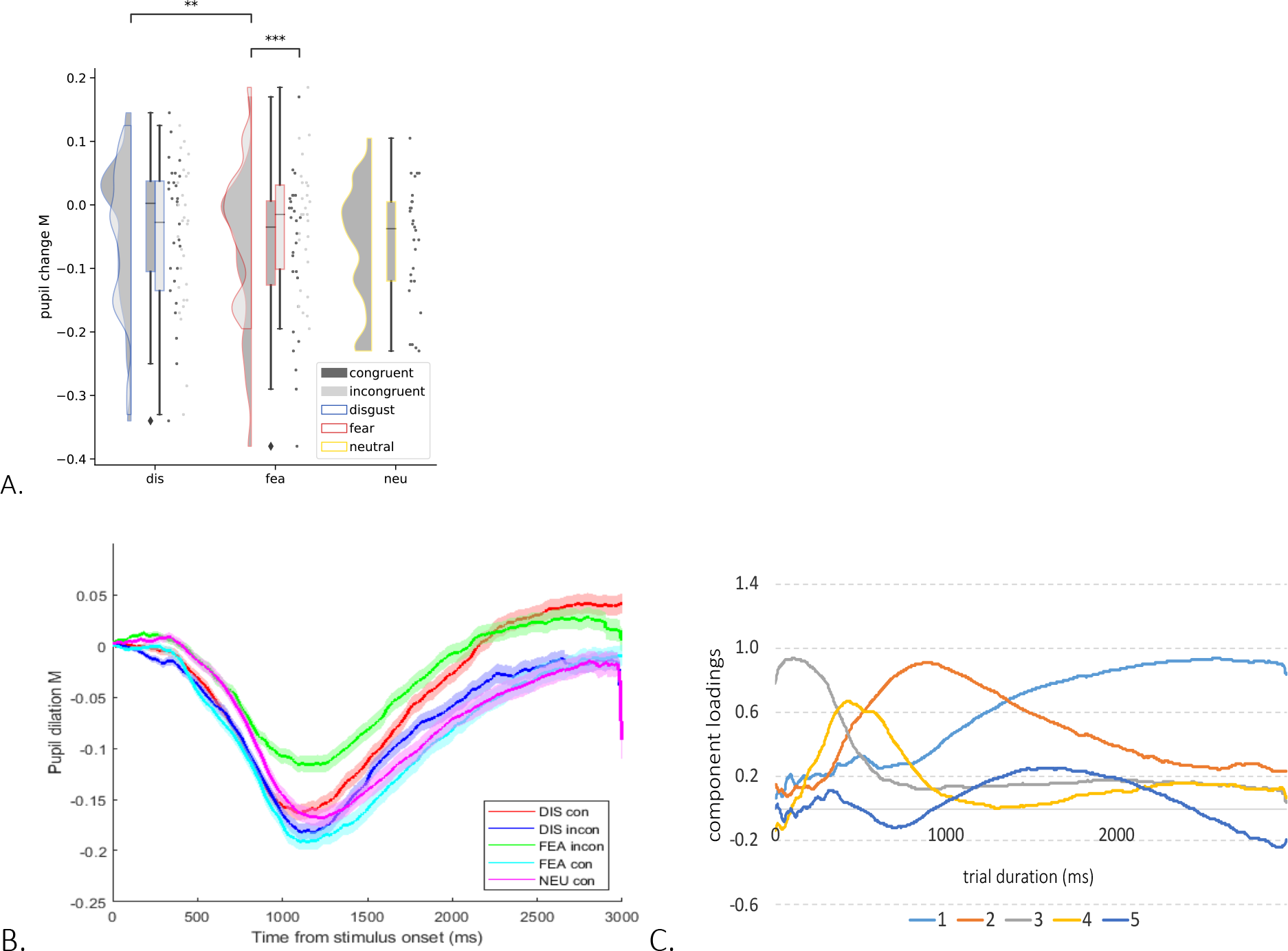
A. Average pupil change during encoding trials (0-3s) is modulated by facial emotion and congruency with emotion schemas. B. Average time course of pupil size during the visual stimulus pair display for correct encoding of faces across emotion and congruency conditions. Modulations by conditions arose over a global light-driven constriction response. C. Temporal PCA on the baseline-corrected pupil change data revealed five pupil components with distinct shapes across time (trial duration of

Previous research suggests that distinct processes can affect pupil size simultaneously, such as visual saliency, task difficulty, cognitive control, or motor responses, which can be dissected by decomposing pupil response through PCA (Granholm & Verney, 2004; Loewenfeld, 1999). In this line, Steinhauer and Hakerem (Steinhauer & Hakerem, 1992) described distinct PCA factors reflecting the contribution of parasympathetic and sympathetic components mixed with other cognitive and visual factors. An early component (750 ms) occurs only with light, presumably reflecting relaxation of iris sphincter muscle (parasympathetic inhibition), while a late component (1750 ms) occurs in darkness and light but increases for emotional stimuli, presumably reflecting constriction of iris dilator muscle (sympathetic activation) (Widmann, Schröger, & Wetzel, 2018).

Here, in an exploratory analysis, we leveraged PCA decomposition to reduce the pupillary response to a subset of factors along the time axis which could account for unique variance in the data (Binetti, Harrison, Coutrot, Johnston, & Mareschal, 2016). PCA is a data-driven approach, completely agnostic about experimental conditions. We found five principal components, characterized by the following temporal features (latencies-to-peak and amount of variance accounted for) after rotation, see Fig. 5C: (1) a late progressive component (peak at 2600 ms; 47.43 % variance); (2) a large intermediate transient component (930 ms, 28.80 % variance); (3) an early rapidly decreasing component (170 ms, 12.15 % variance); (4) a short intermediate component (400 ms, 6.44 % variance); and (5) a slowly varying component (1670 ms, 2.19% variance). These components are consistent with prior work applying PCA to pupil data, including a biphasic pattern (components 2,3,4 vs. 1,5) that may represent separable contributions of the parasympathetic and sympathetic nervous system (Granholm & Verney, 2004; Steinhauer & Hakerem, 1992; Wetzel, Buttelmann, Schieler, & Widmann, 2016; Widmann, Schröger, & Wetzel, 2018). We then computed the degree to which individual loadings on these components during successful encoding was modulated by experimental factors (emotion, congruency and correctness), for all participants, and compared them across conditions. These loadings reflect the degree to which an individual exhibited these distinct temporal characteristics in their pupil response to stimulus pairs.

We found a significant interaction of facial emotions and congruency with emotion schemas for the late progressive component (1) [F(1,29) = 4.854, p = .036, η^2^ = .143] (which did not differ across correctness) and a significant main effect of congruency for a large intermediate transient component (2) [F(1,29) = 7.336, p = .011, η^2^ = .202]. No significant effects were found for the other components. To unpack these results, we ran a post hoc analysis of simple effects comparing different conditions. Paired t-tests for dependent samples showed that the (1) late progressive component, loaded stronger on trials with disgust than fear faces, but only for congruent pairs (p = .006). On the contrary, the (2) earlier transient component, loaded higher on incongruent compared to congruent pairs. This suggests that encoding of disgust faces (when congruent with words) may be associated with higher sympathetic input during slower semantic processing, whereas encoding of incongruent pairs may be more associated with parasympathetic input, temporally aligned with the pupil dilation divergence depending on congruency.

### Correlation with memory performance

Since the above ANOVAs on component loadings found no significant interaction with correctness of subsequent retrieval, we verified whether gaze direction or pupil dilation measures were linked to individual differences in memory performance in an exploratory correlation analysis. Results showed that the more the pupil dilated between 2-3s after trial onset (relative to pre-trial baseline) in response to congruent disgust pairs, the better the subsequent memory for these pairs (r = .444, p = .014). There was no significant relationship between the pupil dilation 2-3s after trial onset and memory performance (r = -.170, p = .359), and these correlation coefficients were significantly different (z = 2.406, p = .008). A similar correlation was found for incongruent disgust pairs (r = .378, p = 036), but not incongruent fear paris, (r =, 051 p = .784), but these correlation coefficients were only marginally different (z = 1.297, p = .097). Likewise, no relationship was found between memory and any of the gaze direction data. These results suggest that only the pupil dilation in a later part of the trial captured individual differences in how various physiological and cognitive processes are related to successful memory formation, specifically for disgust. However, this analysis was exploratory, without any a priori hypotheses, and the results should be treated with caution.

### Effects of emotions and emotion schemas on encoding-related brain activity

In Exp. 2, we used fMRI to measure brain activity associated with successful encoding across different emotion and congruency conditions. We focused our initial analyses on key anatomical regions of interest (ROIs) implicated either in the interplay of memory with emotions and schemas, or in the perceptual processing of faces and words. These regions included the bilateral amygdala (AMY), hippocampus (HC), left inferior frontal gyrus (IFG) and right fusiform gyrus (FG). For each ROI and each condition separately, we extracted activity during successful and unsuccessful encoding of stimulus pairs (i.e. if facial emotion was later correctly or incorrectly retrieved). The engagement of these regions in successful encoding was modulated by both facial emotion and congruency with emotion schemas.

In particular, we found an interaction effect in the left inferior frontal gyrus (IFG) [F(1,17) = 9.029, p = .008, η2 = .033] (Fig. 6A), approaching the correction for multiple comparisons (alpha level of p = .006, tested for each of eight ROIs). Specifically, a higher BOLD signal was found for congruent than incongruent disgust pairs (p = .012), but not fear pairs (p = .143). At the same time, it was higher for pairs including disgust faces if congruent (p = .006) but for pairs including fear faces if incongruent (p = .019). Including the factor of correctness in this analysis resulted in a three-way interaction [F(1,17) = 5.011, p = .039, η2 = .228], which also did not survive a correction for multiple comparisons.

**Fig. 6.**
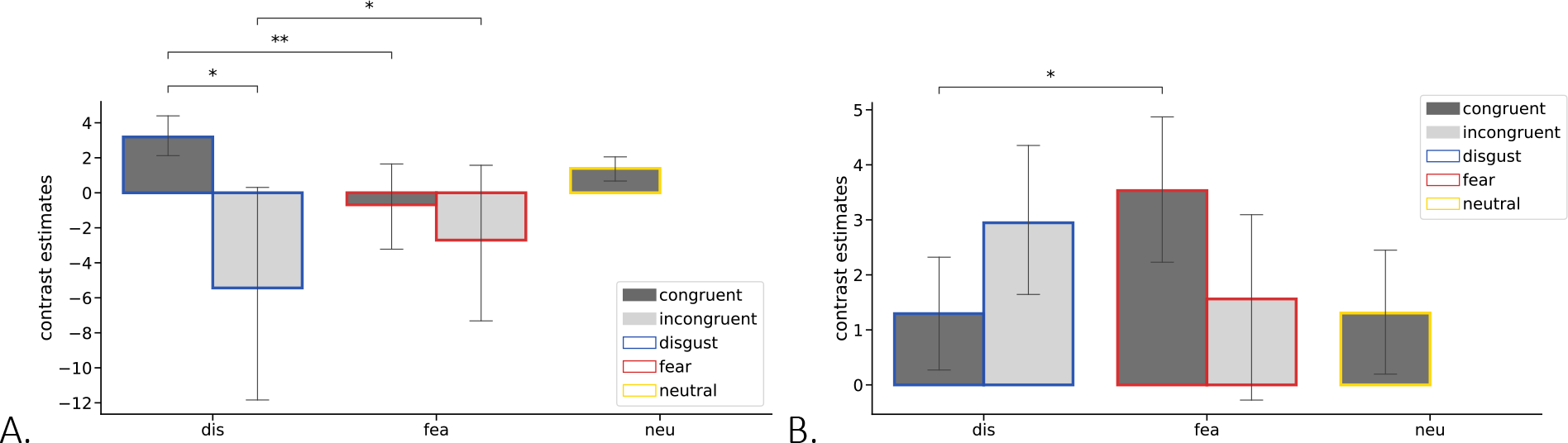
Emotion and congruency effects on the brain activity during successful encoding. Bars represent contrast estimates for successful encoding (followed by correct retrieval) activity in selected ROIs: A. left IFG, B. right FG, split according to emotion and congruency factors (neu condition presented as a point of reference). *Frame colors and labels on x axis indicate the emotion of associated faces to be retrieved: blue = disgust, red = fear, yellow = neutral; bar colors indicate the congruency with emotion schemas: dark grey – congruent, light grey – incongruent. Error bars represent one standard deviation; DIS – disgust, FEA – fear; * p < .05, ** p < .01*.

Likewise, when considering only old pairs congruent with emotion schemas we found an interaction effect between emotion and correctness [F(2,34) = 4.286, p = .039, η2 = .201], such that it was more activated during encoding of disgust- than fear-related (p = .019) and neutral (p = .013) pairs, but only if successfully encoded. However, this effect did not survive a correction for multiple comparisons. This result suggests that the role of IFG during successful encoding was particularly strong for pairs congruent with emotion schema, but specifically for disgust, possibly supporting more semantic processing.

An interaction of facial emotion by schema congruency and correctness was also found in the right FG [F(1,17) = 6.198, p = .023, η2 = .267] (Fig. 6B). Contrary to the left IFG, it was driven by the higher right FG activation to congruent fear compared to congruent disgust (p = .027) among correctly encoded pairs. When comparing between emotion categories and neutral among congruent pairs only, we also found an interaction with correctness [F(2,34) = 4.385, p = .030, η2 = .205], but none of these effects in FG survived a correction for multiple comparisons. This result suggests an important role of right FG during encoding of congruent pairs related specifically to fear, suggesting more sensory processing of faces. No significant results were found in other ROIs, all reported in supplementary materials (Fig. S5)

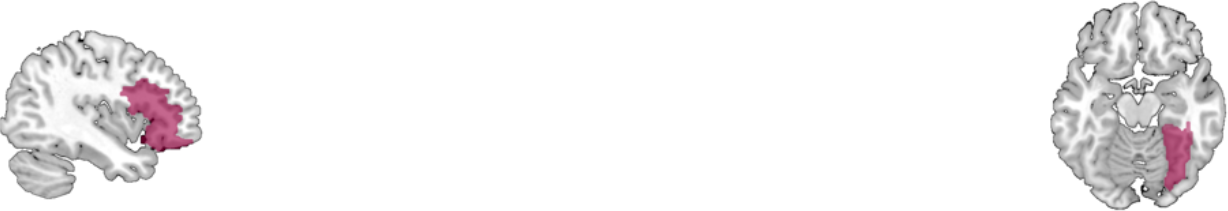

To complement our ROI analyses based on a priori hypotheses about memory modulations, we conducted a whole-brain random-effect analysis. Specifically, we computed a linear contrast testing for a positive interaction between emotion and congruency during successful encoding, masked by exclusion of the same interaction contrast for unsuccessful encoding. This masking procedure ensures that we obtain significant brain activation for successful encoding. The results showed a higher BOLD signal in left IFG and medial prefrontal cortex, together with parietal regions, precuneus and FG (Fig. 7a and 7b, all statistics presented in Supplementary Table S3). These results accord with previously reported effects of schemas and semantic congruency on memory, but in addition reveal a differential modulated by the emotional category of facial expressions.

**Fig. 7.**
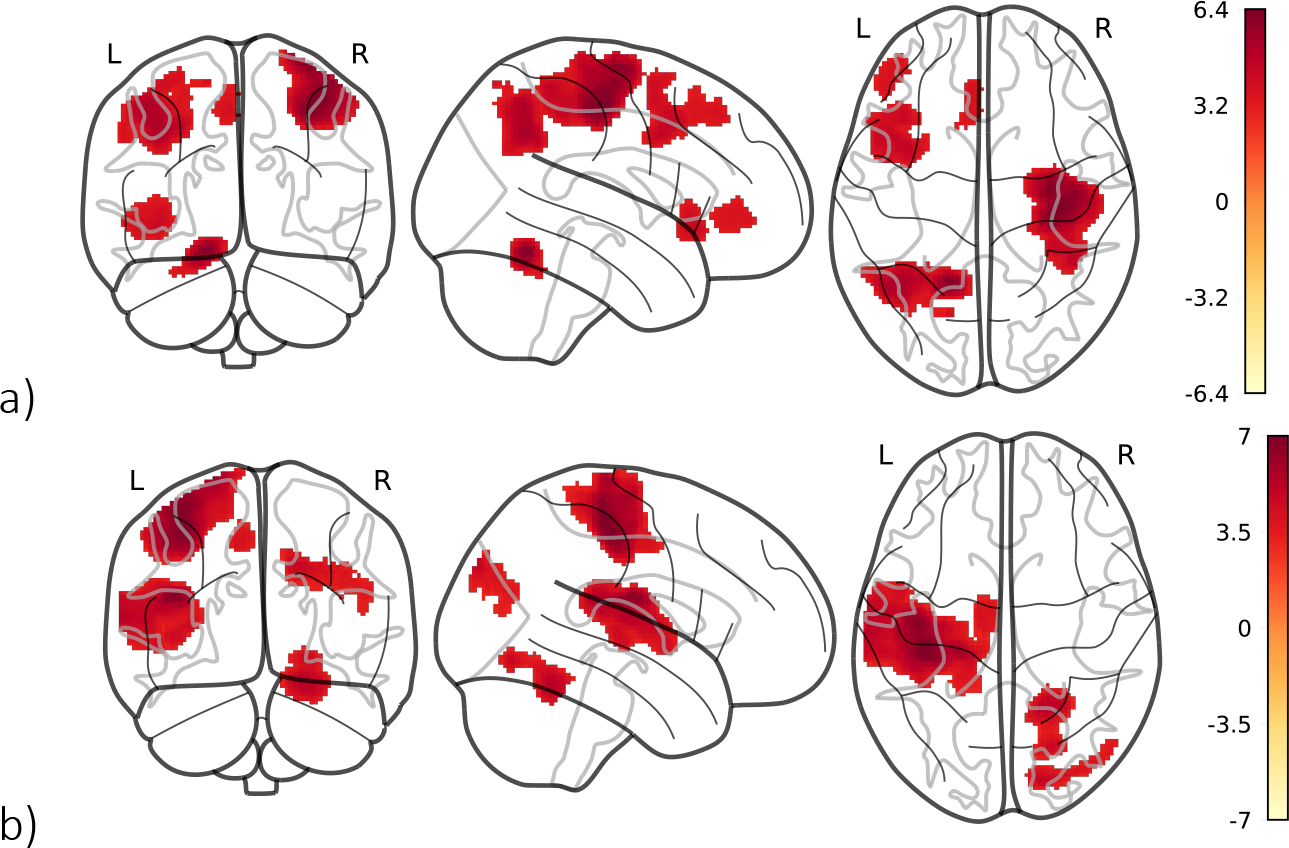
Brain activations showing interaction effects between emotion and congruency during successful encoding (exclusively masked by the same interaction during unsuccessful encoding) in two possible directions: a) activations higher for disgust > fear and congruent > incongruent; (a) activations higher for the reverse contrasts. Both a) and b) contribute to the interaction effect. *Colour scale bars represent a range of t-values; voxel-wise height threshold of p < 0.001 (unc.) combined with a Monte Carlo simulation cluster-level extent threshold (nn = 2, two-sided, p = .001, alpha = .05) of k = 172 voxels. The glass brain depiction complemented with specific brain slices, Fig. S7*.

## Discussion

In the present study, we combined behavioural measures of retrieval with eye-tracking and fMRI to investigate unresolved issues about the interplay between schema and emotion effects on human memory. Previous behavioural studies have consistently reported memory enhancement for associations congruent with our prior knowledge of the world, or schemas (Bein et al., 2015b; DeWitt et al., 2012; Kole & Healy, 2007). Recent neuroimaging studies have begun to point to the underlying substrates of these processes (Gilboa & Marlatte, 2017; Preston & Eichenbaum, 2013b). For example, prior knowledge promotes separation of similar memories in the hippocampus, but also assimilation of newly learnt associations into existing structure of knowledge in the left inferior frontal gyrus (Bein et al., 2020). However, crucial question remains open - how these effects interact with other mnemonic influences such as those due to the emotional content of events, and whether emotional congruency with context could also serve as a schema to facilitate memory formation and retrieval.

Using a novel face-word pair association paradigm and a multi-method approach, we showed across two independent studies with large samples of participants, in a situation resembling real-life communication, that emotion category (disgust vs. fear) and congruency with emotion schema (congruent vs. incongruent) both affected associative memory and related physiological and neural mechanisms. Overall, emotional congruency of words and faces facilitated memory for these associations. Behaviourally, this result was reflected by the correctness of retrieval, as well as by the reaction times of retrieval, showing that memories were more accessible and a mnemonic decision was easier to make (Brod, Werkle-Bergner, & Lee Shing, 2013). Physiologically, on the contrary, encoding of incongruent vs. congruent pairs loaded higher the early transient pupil component reflecting a parasympathetic input. At the neural level, we observed an engagement of the left IFG during successful encoding of emotionally congruent vs. incongruent disgust-related pairs, a region previously implicated in the effects of prior knowledge and mental schemas in memory (Bein, Reggev, & Maril, 2014a; Bein et al., 2020). These results suggest that emotional congruency may indeed affect memory in a similar way to semantic congruency, typically examined in previous studies (Bein et al., 2015b), and could thus serve as another sort of schema. This finding accords with situations of real-life communication where it is more probable to encounter facial expressions and verbal messages expressing the same emotion (unlike, for instance, in the case of irony (Banasik-Jemielniak, 2021; Nuber, Jacob, Kreifelts, Martinelli, & Wildgruber, 2018)).

Our results indicate that emotional congruency effects differed between emotion categories of disgust and fear. At the behavioural level, the memory facilitation by congruency was weaker for fear- than disgust-related stimuli, possibly driven by generally lower associative memory due to fear-related information (found when compared to neutral pairs across congruent conditions). This finding is in line with a large body of research indicating that negative affect (typically fear in the previous studies) impairs associative memory (James A. Bisby & Burgess, 2014). The impaired memory for associations was previously demonstrated both for when a negative event element was encoded close in space (i.e. presented simultaneously), and in time (i.e. presented sequentially, negative first (James A. Bisby, Horner, Bush, & Burgess, 2018)). Such effect has been related to a disruptive role of amygdala (J. A. Bisby & Burgess, 2017; James A. Bisby, Horner, Horlyck, & Burgess, 2016; Madan, Fujiwara, Caplan, & Sommer, 2017; Murray & Kensinger, 2014; Okada et al., 2011) and an increase in functional connectivity between the hippocampus and amygdala when encoding associations with negative compared to neutral stimuli (Dandolo & Schwabe, 2018). Of note, modulation of contextual memory by fearful or disgusted faces was recently questioned due to a n unsuccessful replication attempt (H. A. Chapman, 2020), yet the paradigm differed from ours in multiple aspects including the sequential vs. simultaneous encoding. Another study also reported that, during encoding, participants made substantially fewer saccades between paired negative pictures than between neutral pictures, which was related to worse associative memory (Madan et al., 2017). Here we found a similar drop in gaze switches (or saccades) between words and faces for congruent fear-related pairs compared to the disgust condition, as well as for incongruent compared to congruent disgust, which was in line with differences in retrieval rate.

It could be that these emotion-specific effects on memory for congruent pairs were due to higher arousal. Because our encoding task prioritized the memory of faces over the memory of words, pairs including fearful faces possibly gained higher saliency and were more arousing, consistent with the observed pupil change pattern, thought to track levels of locus coeruleus noradrenaline activity (Clewett et al., 2018). This in turn might lead to better subsequent memory for these faces, with no effect on lower-priority word cues. Such effects can be triggered by an increased neural gain (Aston-Jones & Cohen, 2005a; Eldar, Cohen, & Niv, 2013), a process by which patterns of high neural activation are further excited, whereas patterns of low neural activation are further inhibited, which in turn influences learning and memory (Eldar et al., 2013). However, prioritization of memory is typically induced in a more powerful way with other paradigms such as a monetary incentive encoding task, whereas in our case this would emerge only indirectly through strategic use of our instruction to memorize facial emotions. Moreover, in our task, the retrieval of targets (faces) was cued by words and we cannot disentangle the contribution of both to successful memory of associations. Therefore, further experiments are necessary to verify the role of such implicit saliency gain and noradrenergic system effects on memory performance. Furthermore, the only link between the pupil response (2-3s after trial onset) and memory we observed only for disgust-related faces (congruent and incongruent), which speaks against the hypothesis of arousal driving a reduced memory for fear-related context.

Another possible explanation of our results derives from the emotion and congruency interaction effects observed in memory performance and concomitant activation of left IFG. This brain region is implicated not only in memory schemas, but also in semantic processing. Several studies have highlighted the role of IFG in either the retrieval or selection of semantic knowledge (Thompson-Schill, D’Esposito, Aguirre, & Farah, 1997; Wagner, Paré-Blagoev, Clark, & Poldrack, 2001). Accordingly, some authors suggested that event congruency enhances episodic memory encoding through semantic elaboration and relational binding (Bernhard P. Staresina, Gray, & Davachi, 2009). Our results show that the IFG activity was higher during encoding of congruent vs. incongruent pairs, but only in the case of disgust, suggestive of more semantic processing involved in the learning of disgust-related associations. This interpretation was supported by the gaze direction results showing more fixations on words, and more gaze switches between words and faces for congruent disgust compared to fear. Moreover, we found more variance in pupil dilation explained by a late progressive component (related to the sympathetic activation) for congruent disgust than for fear-related pairs, which is in line with slower semantic processing.

On the other hand, we observed that during encoding of congruent fearful pairs, participants spent more time observing faces than words, associated with enhanced sensory processing in the right FG. This accords with existing evidence for involuntary attraction of attention by threat-related stimuli (Furtak et al., 2020), especially fearful faces (Vuilleumier, Armony, Driver, & Dolan, 2001). The latter study also demonstrated that the manipulation of attention strongly modulates the FG (but not amygdala) response to faces, which partly aligns with our results. A qualitative drop in FG engagement for incongruent vs. congruent fearful pairs may reflect an additional division of attention, accompanied by a significant decrease in pupil dilation and time spent observing faces, but a longer time spent observing words. Conversely, a qualitatively greater FG engagement to incongruent vs. congruent disgust-related pairs might reflect the attraction of attention to fearful words paired with disgusted faces, paralleled with a significant increase in gaze switches.

It should be noted that contrary to existing literature, we found a limited relationship between our eye-tracking and pupillometry results and memory performance, paralleled with the engagement of MTL at the neural level. Contrary to former studies, we did not observe memory-related differences in gaze patterns during encoding (Hannula et al., 2010; Hannula, Ryan, Tranel, & Cohen, 2007; Ryan & Shen, 2020). It was previously shown that successful learning of associations should be reflected by longer and more frequent fixations, as well as more gaze switches between memorized information (Eckstein et al., 2017; Madan et al., 2017). Here, we observed an interaction between emotion and congruency on the fixations on words, faces and gaze switches between the two, yet no across-subject correlation with memory performance. However, previous evidence was based on experimental paradigms using mostly neutral, congruent associations, and may not be evident on top of the complex schema and emotion effects we observed. Likewise, previous research suggested that pupil size can reflect a successful encoding (Kucewicz et al., 2018; Naber, Frässle, Rutishauser, & Einhäuser, 2013; Võ et al., 2008), in line with the results of our exploratory analysis (specific to disgust and 2-3s after trial onset). We did not find a link between memory and different temporal components of pupil response that were modulated by emotion and congruency with emotion schemas. However, previous literature suggested only a link with temporal memory (Clewett, Gasser, & Davachi, 2020b), not tested in the present study.

To conclude, our results elucidate a previously unknown pattern of memory modulation by emotional congruency, acting as another type of mental schema to facilitate memory formation. In addition, we show this effect is differentially expressed according to distinct emotion categories, showing another level of emotion-specific effects in human memory formation. These parallel influences presumably reflect different mechanisms of learning associations, with more semantic processing for disgust and more attentional effects for fear.

## Acknowledgements

This work was funded by the National Science Centre Poland (2015/19/N/HS6/02376), MR was also supported by the Polish Ministry of Science and Higher Education (1625/MOB/V/2017/0). We would like to thank: Dawid Droździel and Bartosz Kossowski for their technical support with the MR scanner, Remi Neveau for his help with eye-tracking data analysis, Maria Kulesza for her help with eye-tracking data collection, Lila Davachi and David Clewett for their inspiring advice on the pupil PCA, Agata Bochyńska for her advice on eye-tracking and pupillometry data analyses, Oded Bein and Tarek Amer for their useful conceptual comments on the manuscript.

## Data availability statement

Norms for words used in the experiment are available as supplementary materials to the previous publications: https://lobi.nencki.gov.pl/research/18/

All data analysed and presented in the current manuscript are publicly available: https://osf.io/y9n2k/?view_only=b19f39928f194ae19716b98c99745801

## Ethics statement

In the present study, all participants provided written informed consent and were financially compensated. The local Research Ethics Committee at Faculty of Psychology, University of Warsaw approved the experimental protocol of the study.

## Author contributions

Monika Riegel – conceptualization, methodology, Investigation, data curation, resources, software, formal analysis, visualization, validation, writing - original draft, writing – review & editing, project administration, funding acquisition

Marek Wypych – validation, data curation, software, formal analysis

Małgorzata Wierzba – investigation, data curation, resources, software, formal analysis, validation, writing – review & editing

Michał Szczepanik – software, data curation, validation, formal analysis

Katarzyna Jednoróg – methodology, writing - review & editing

Patrik Vuilleumier – conceptualization, methodology, formal analysis, supervision, writing - review & editing

Artur Marchewka – conceptualization, methodology, resources, supervision, writing - review & editing, funding acquisition

## Supplementary materials

**Table S1.**
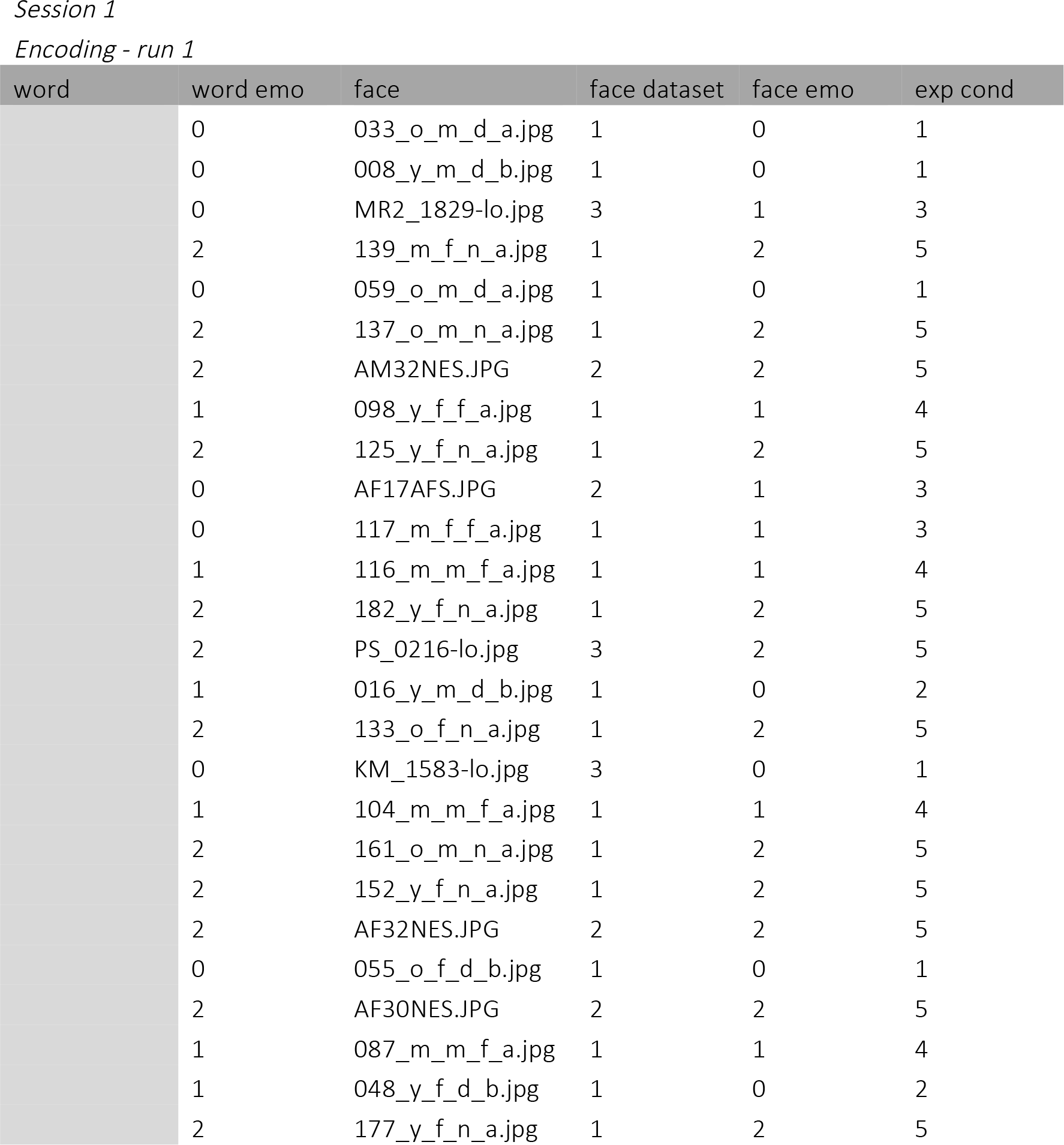
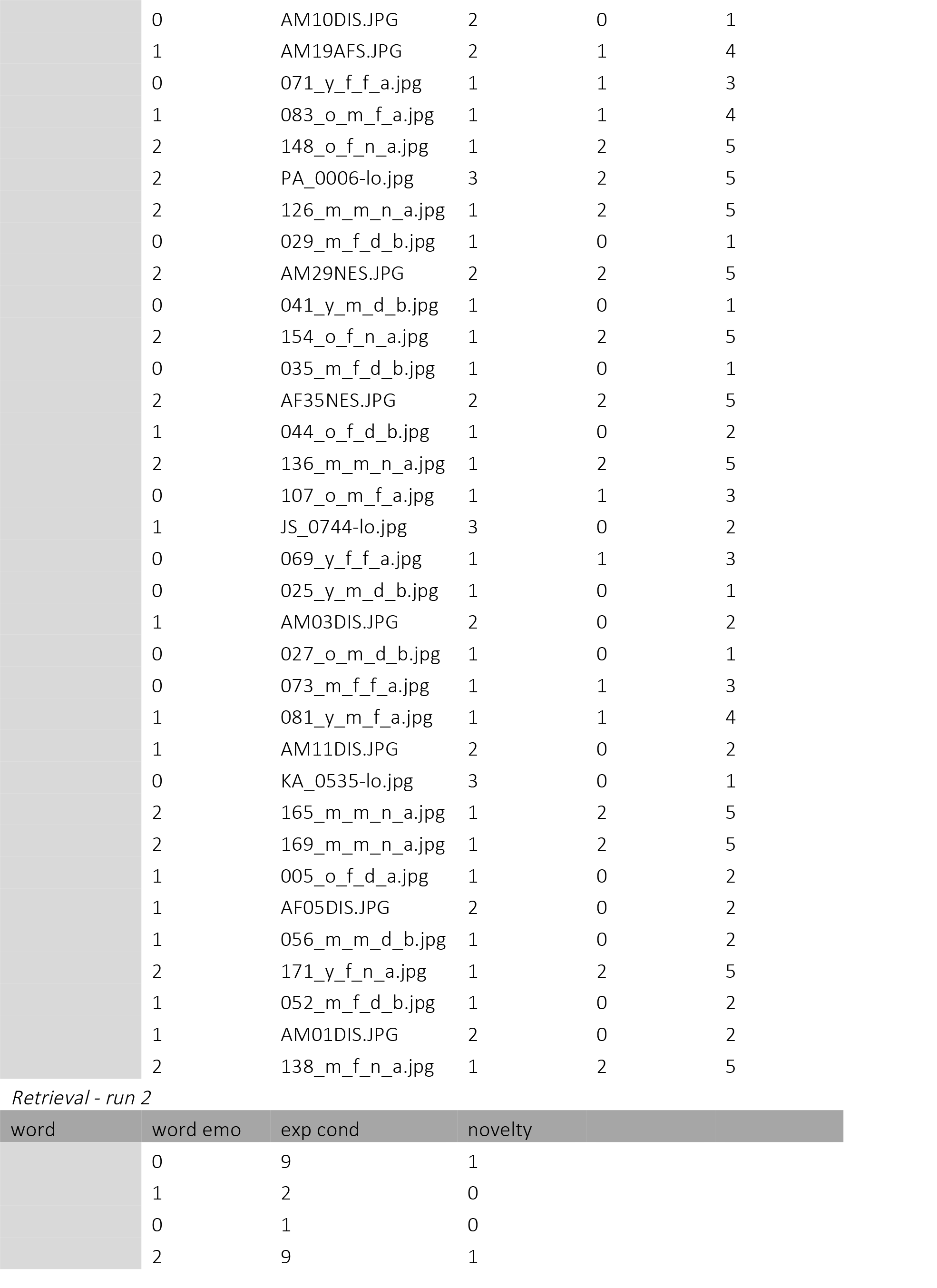
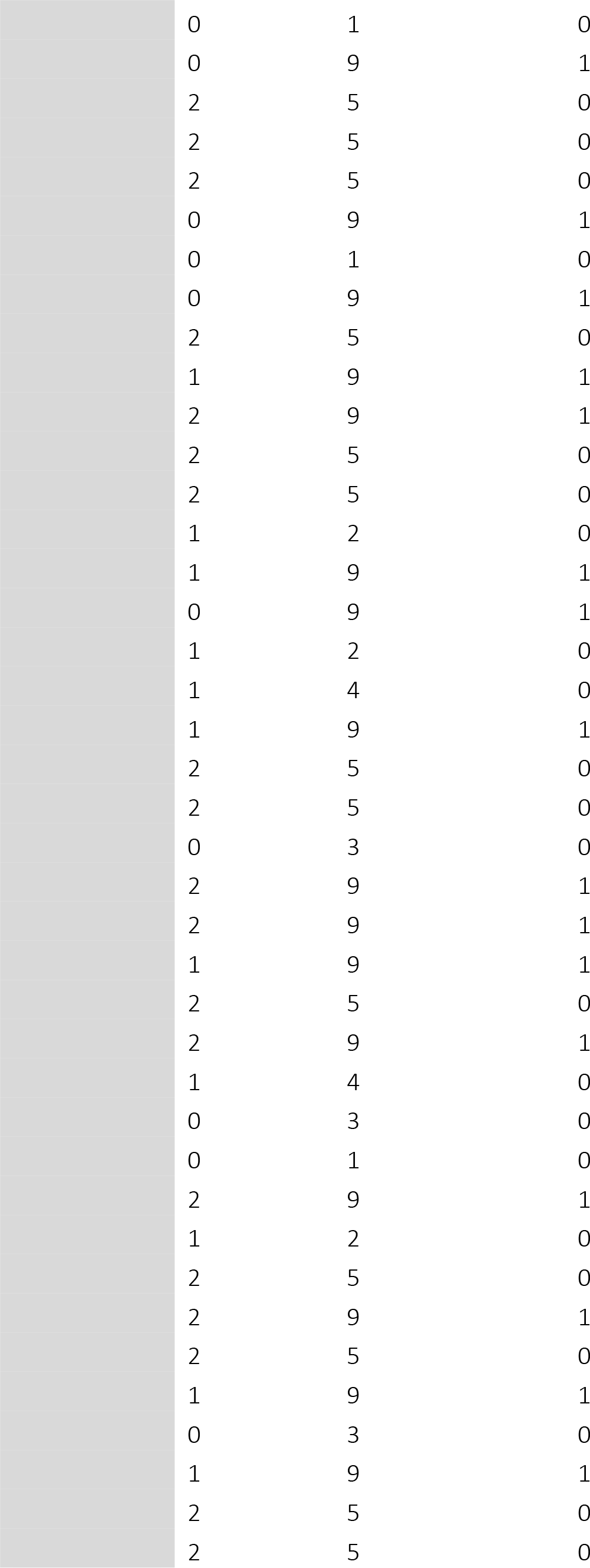
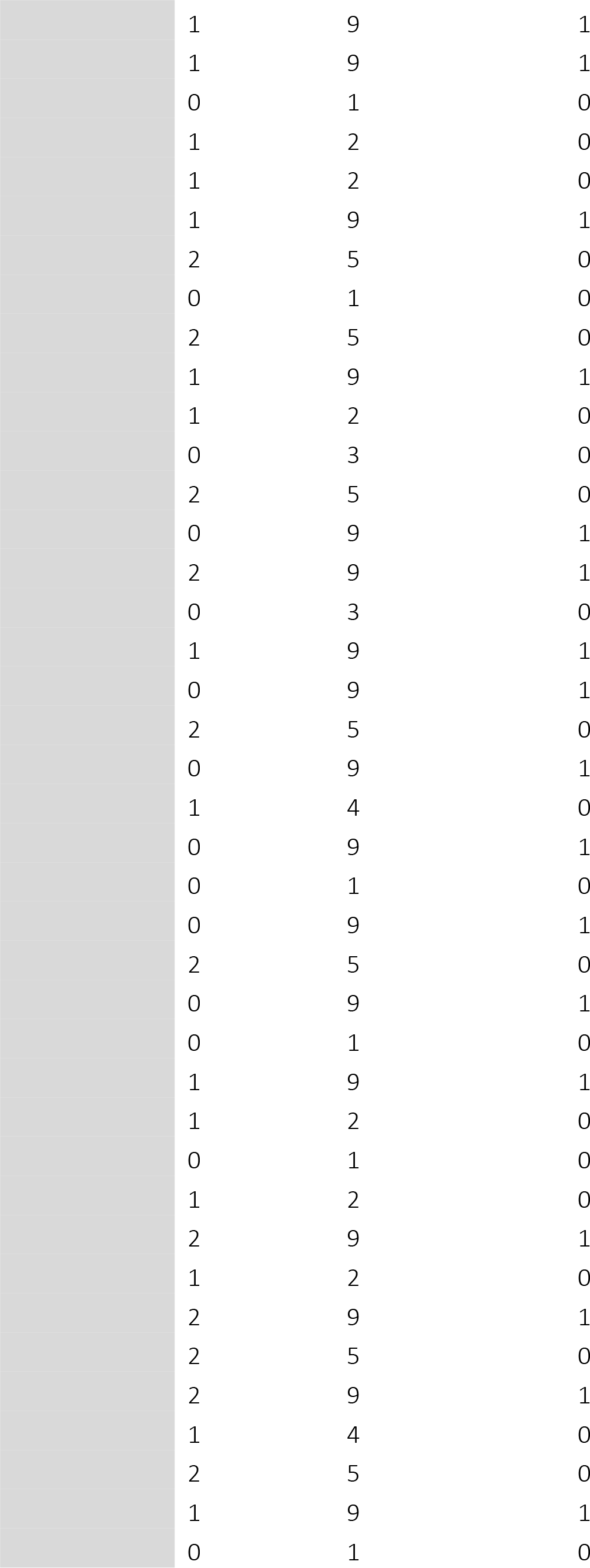
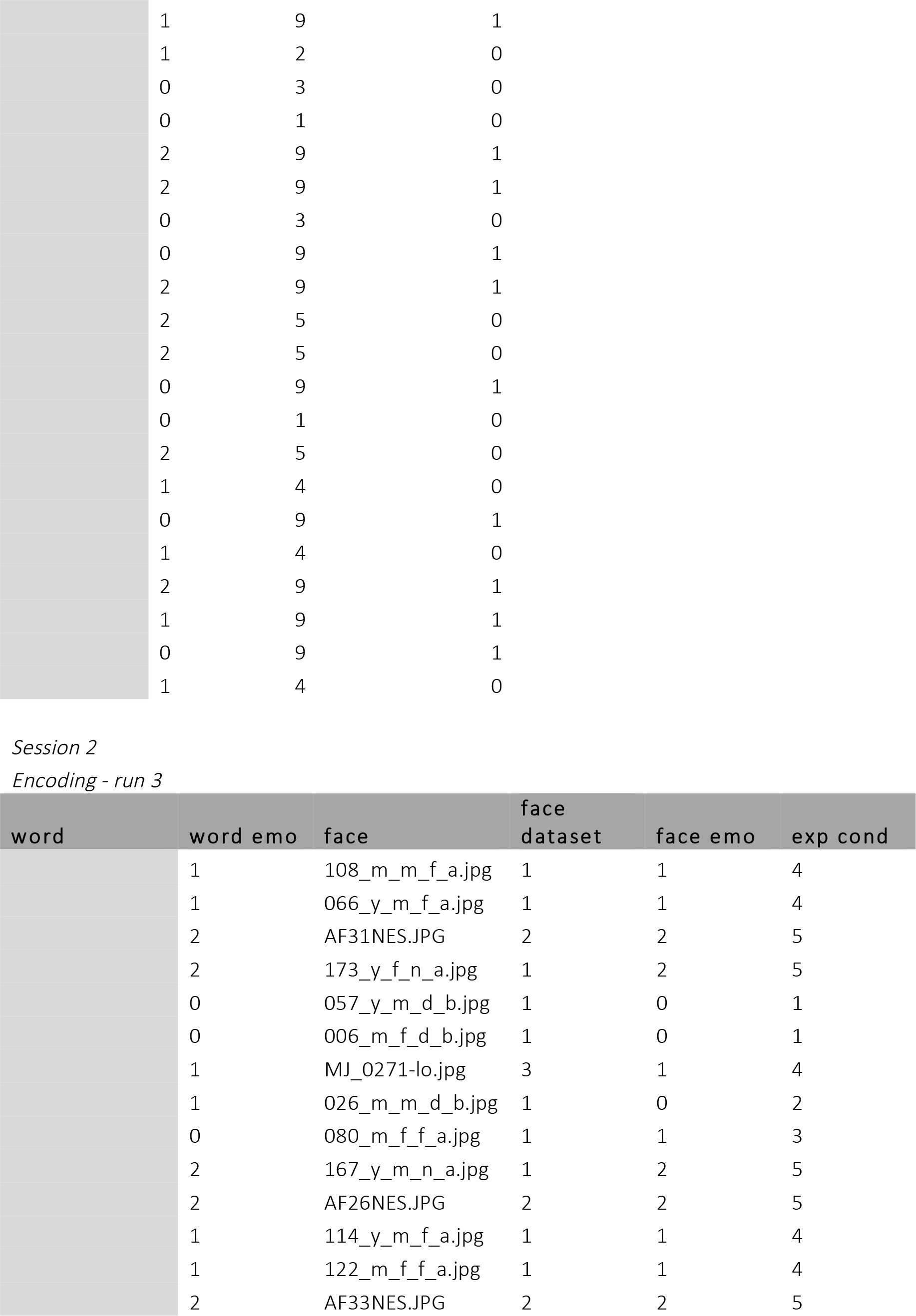
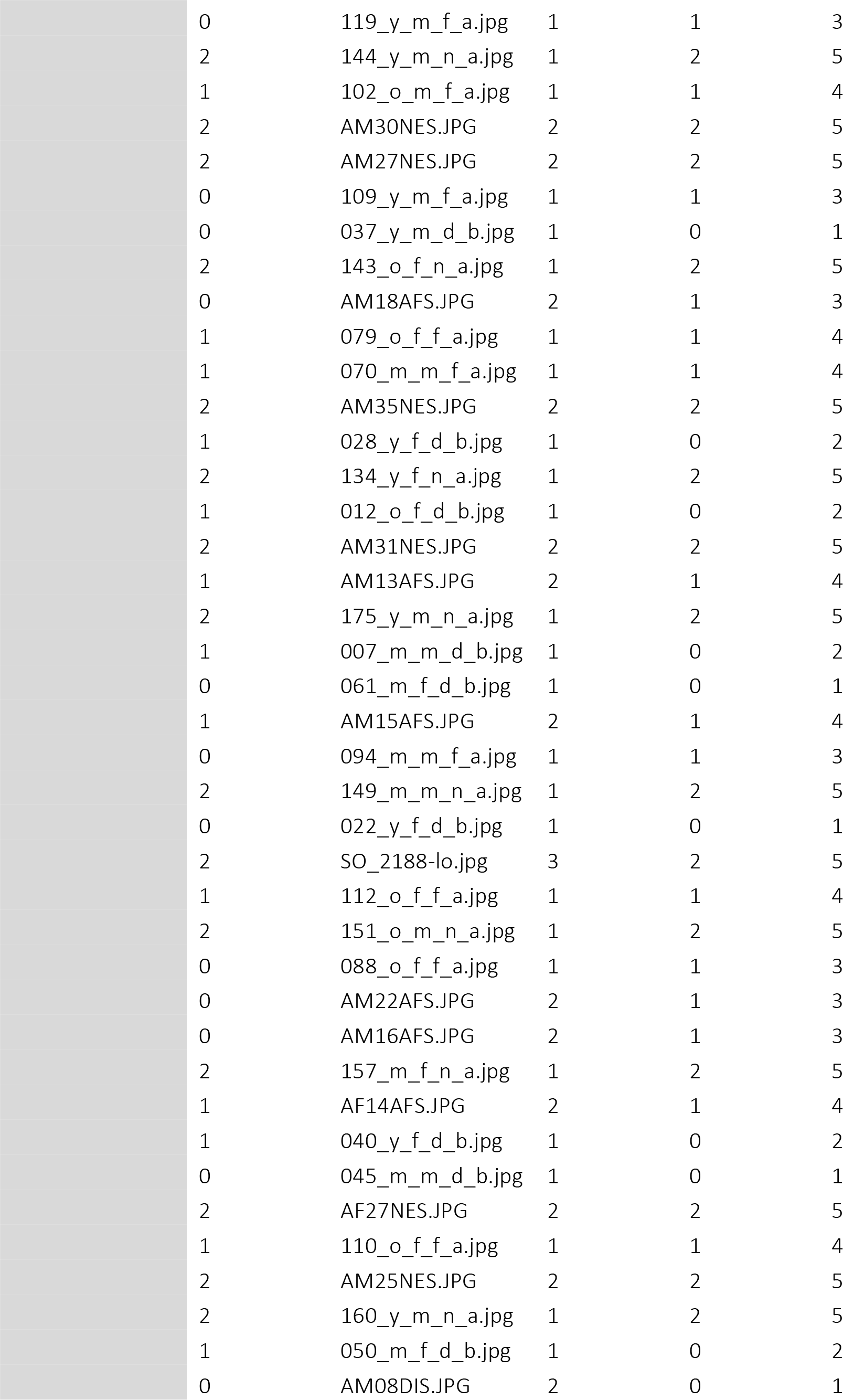
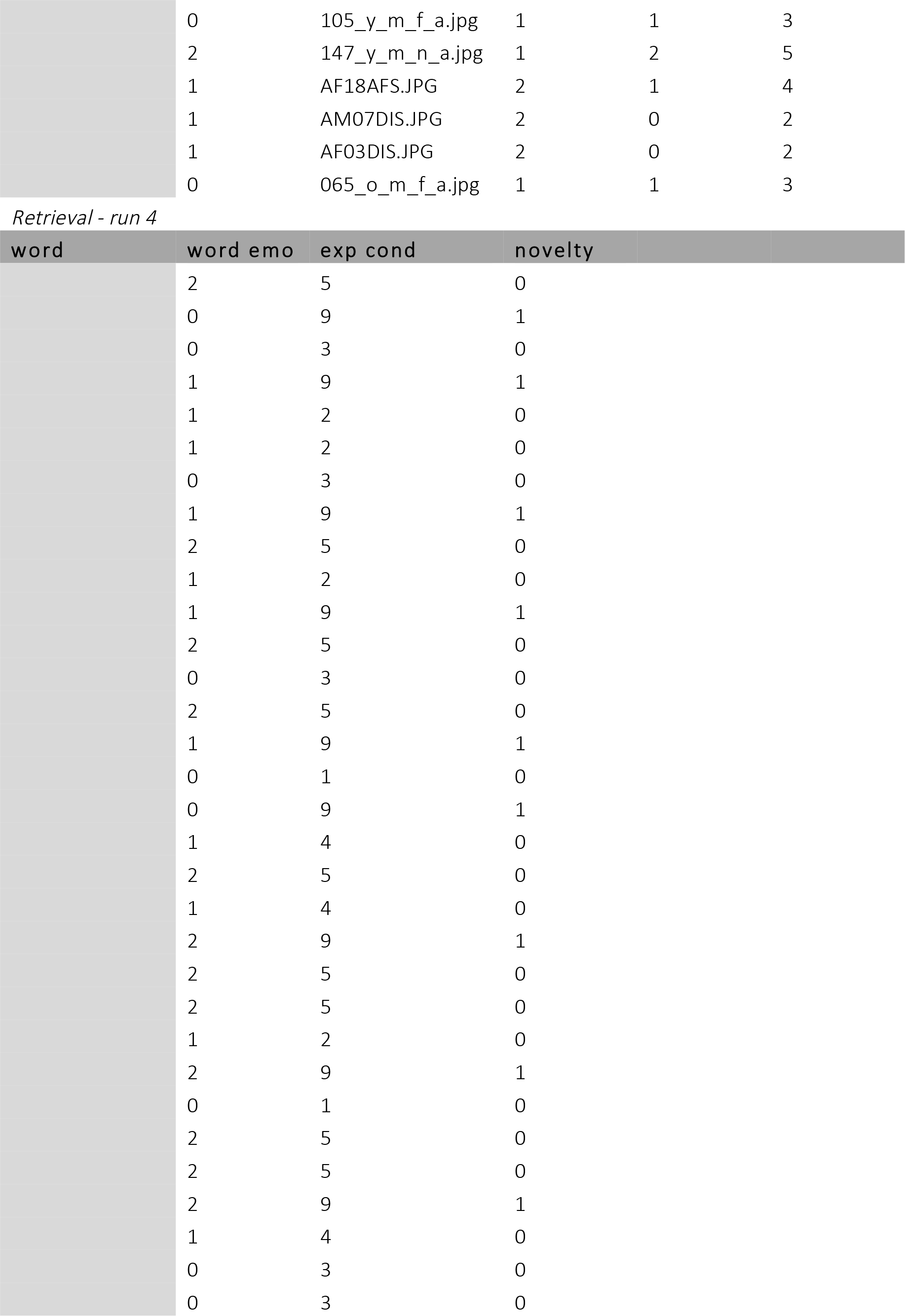
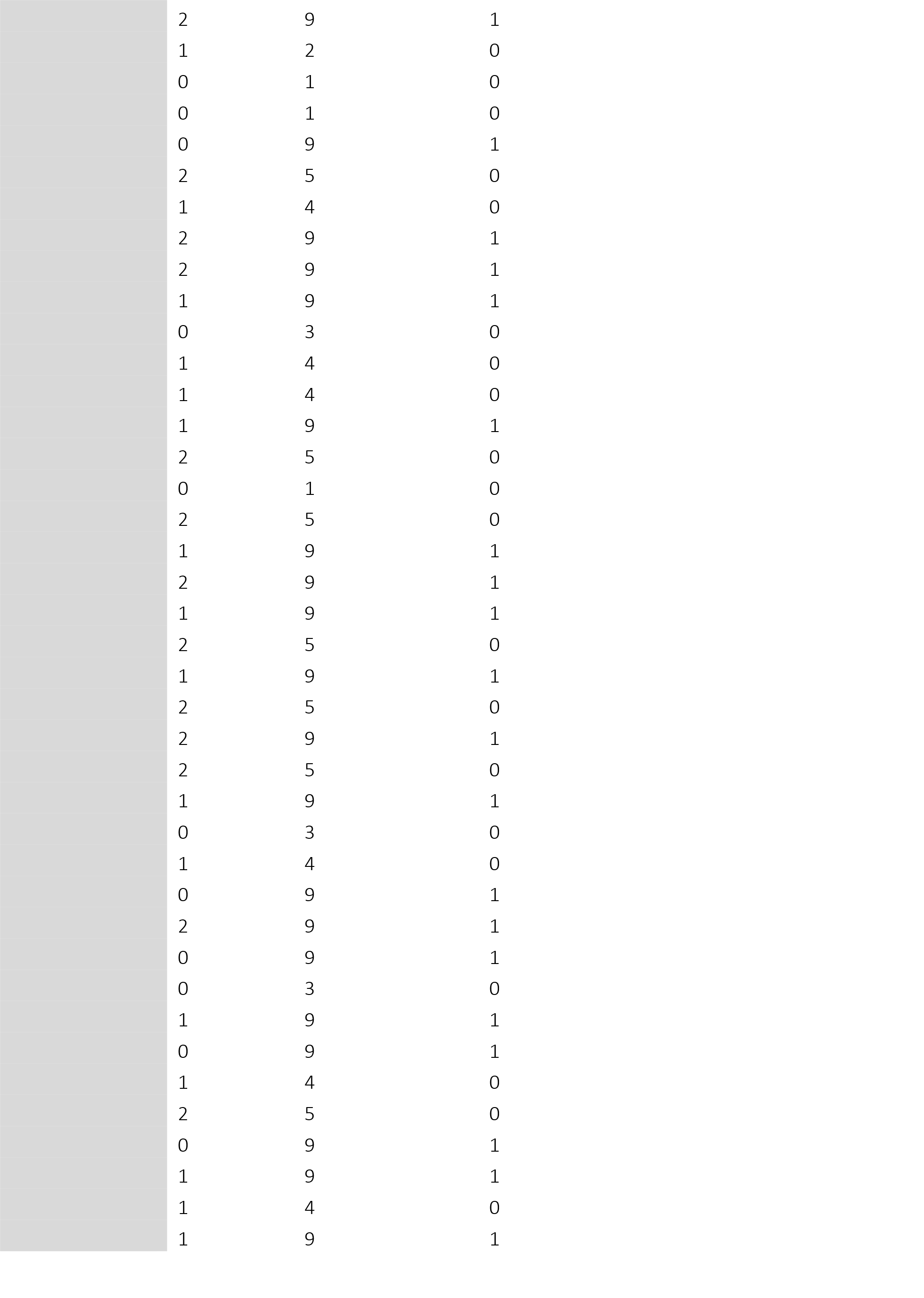
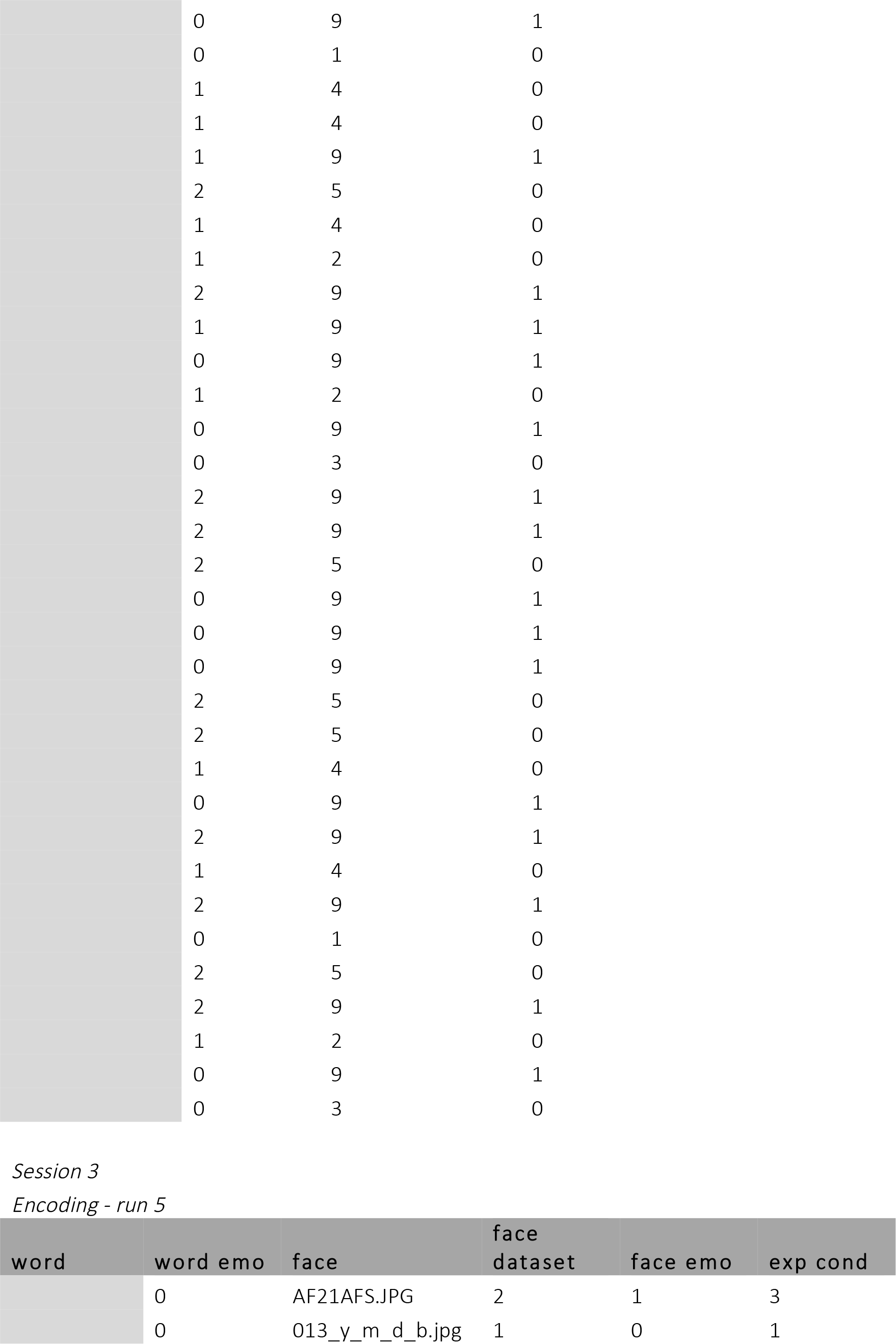
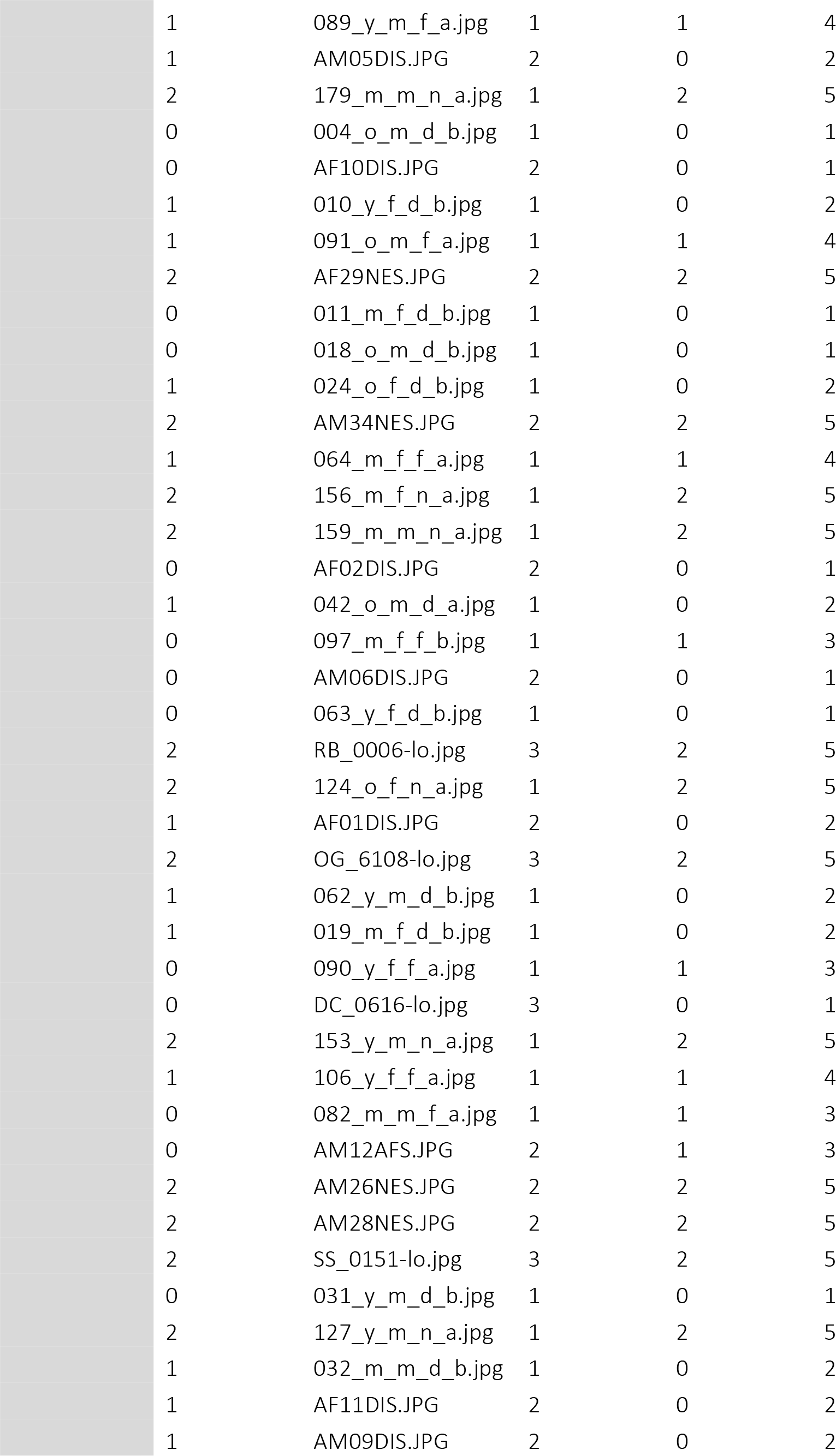
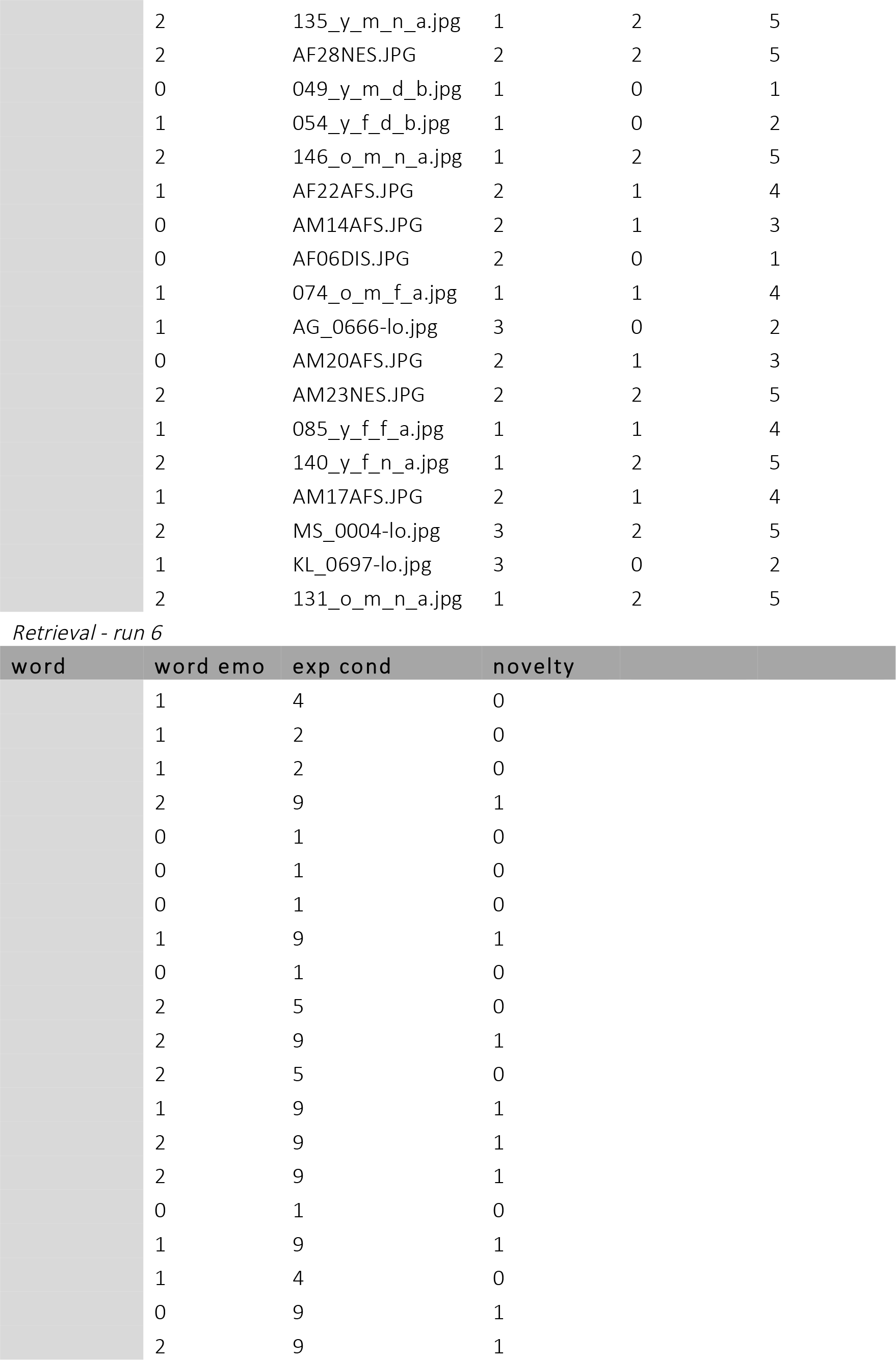
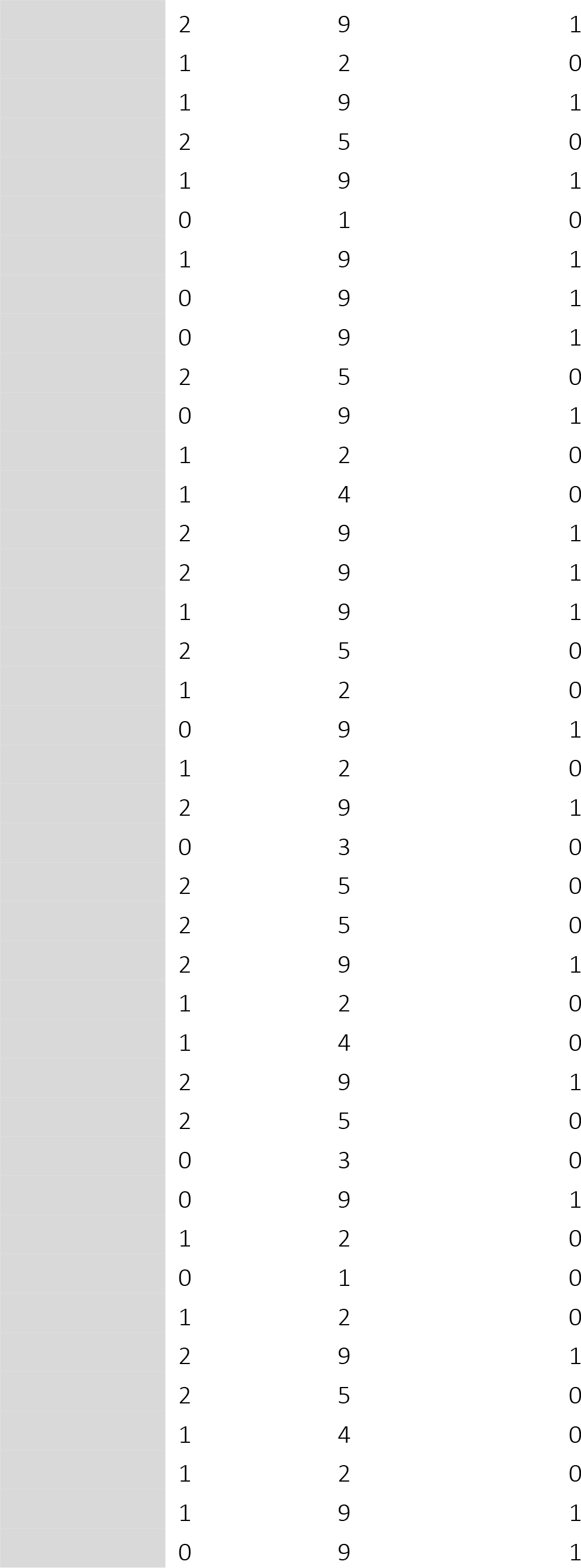
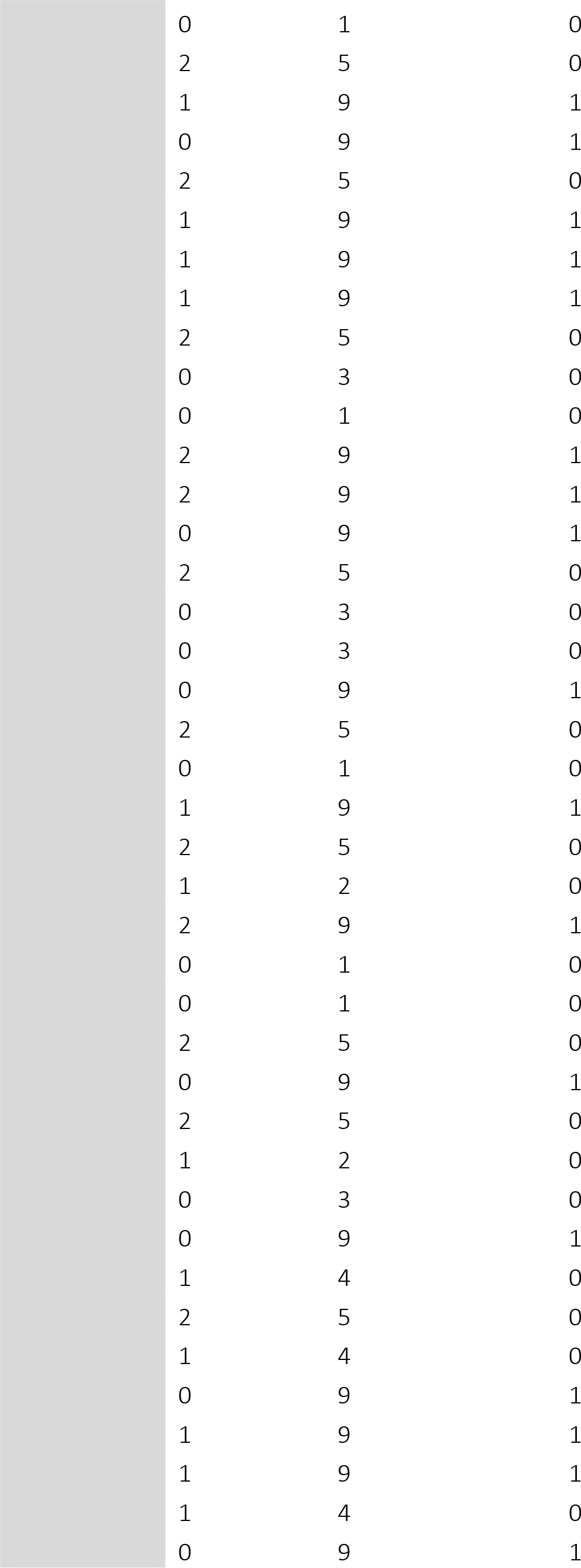
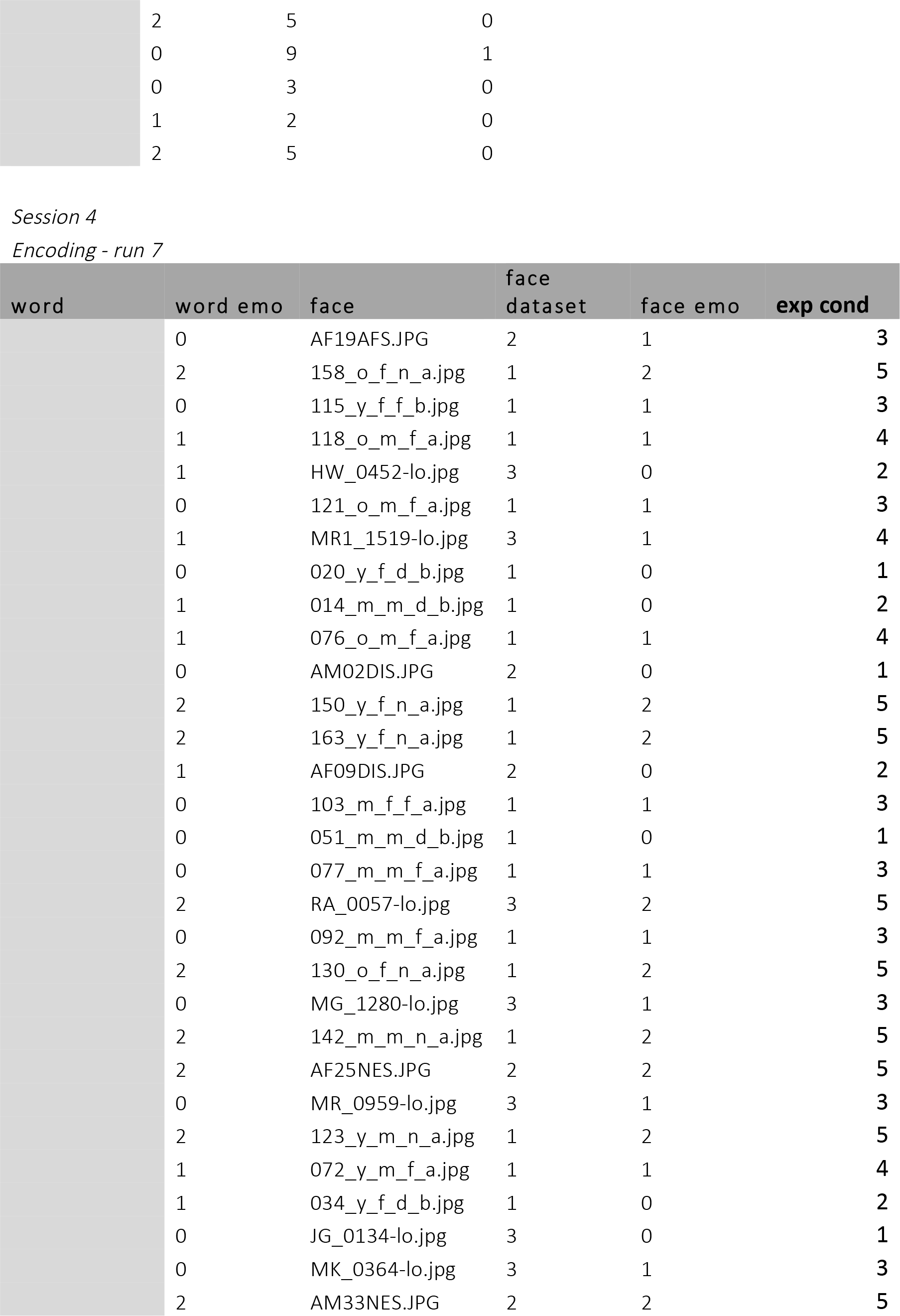
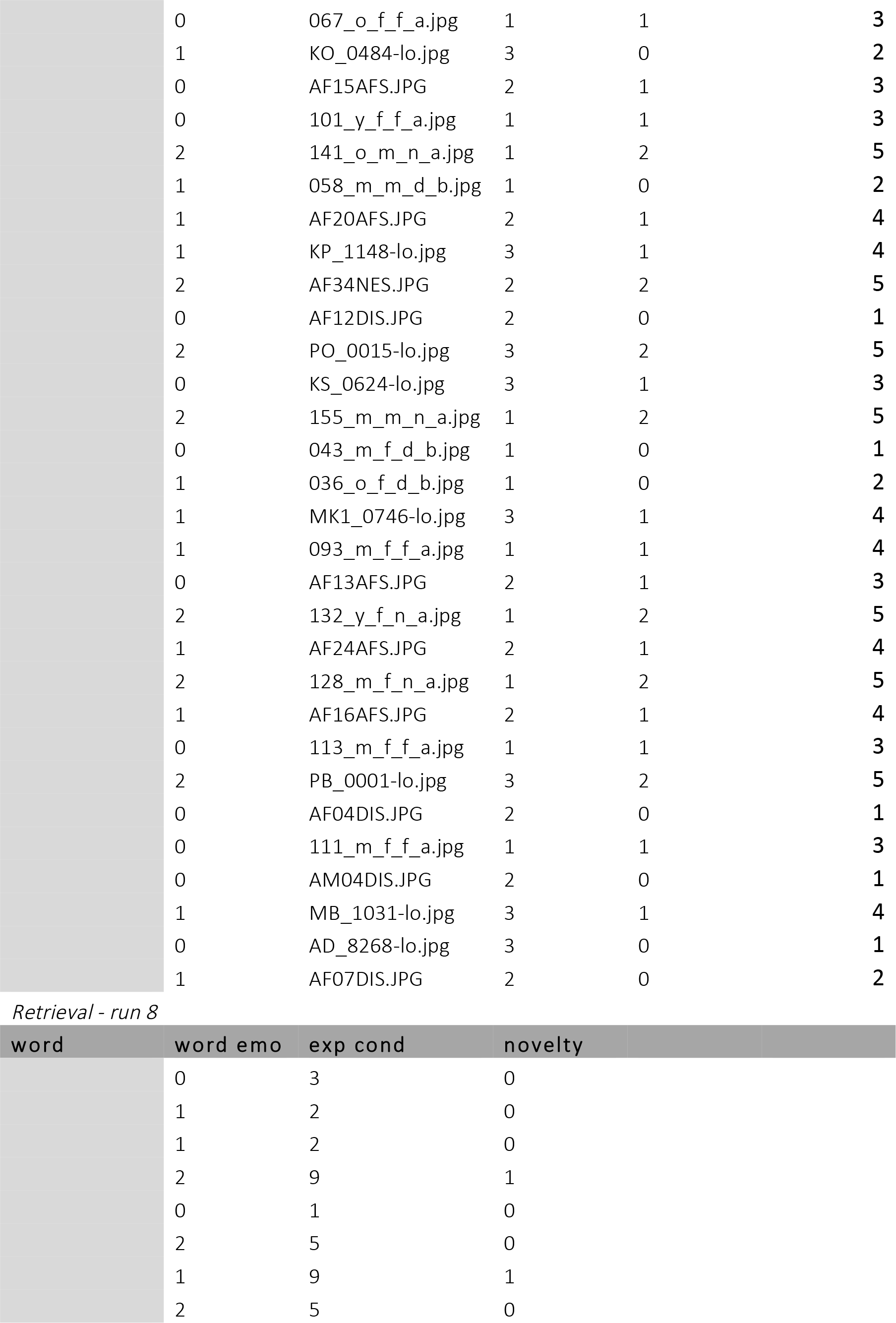
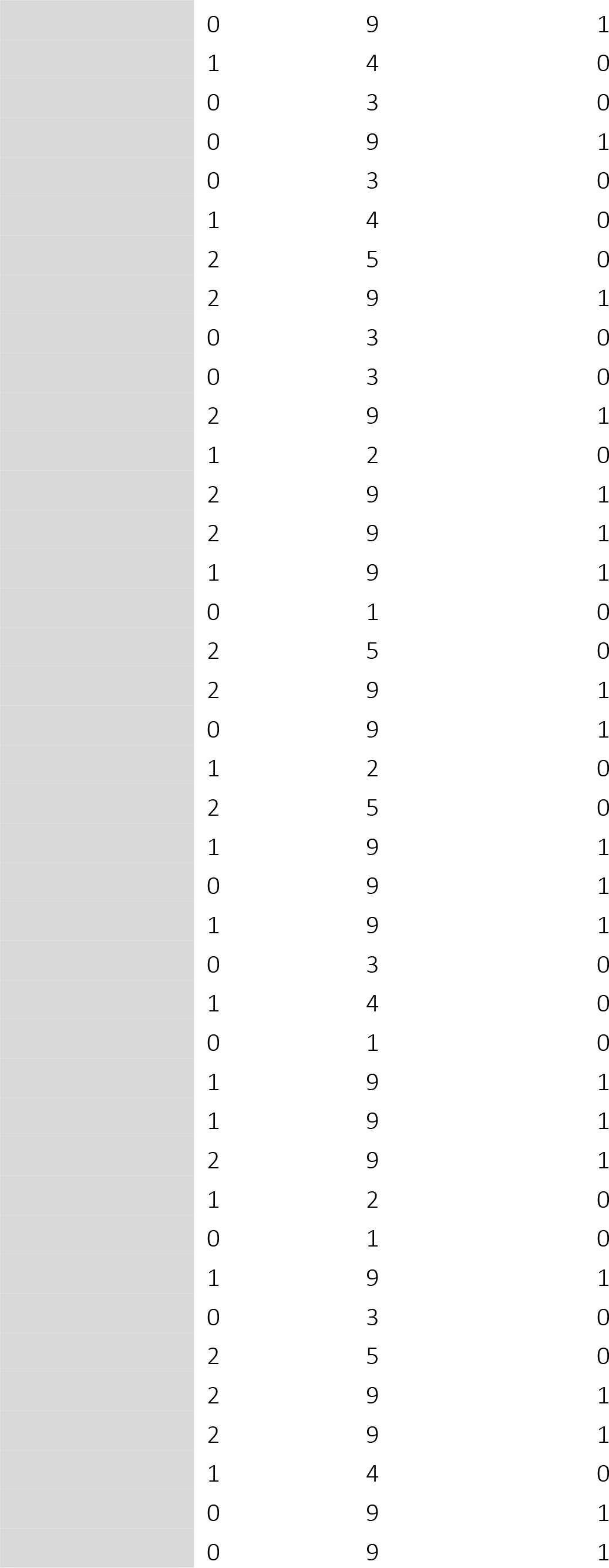
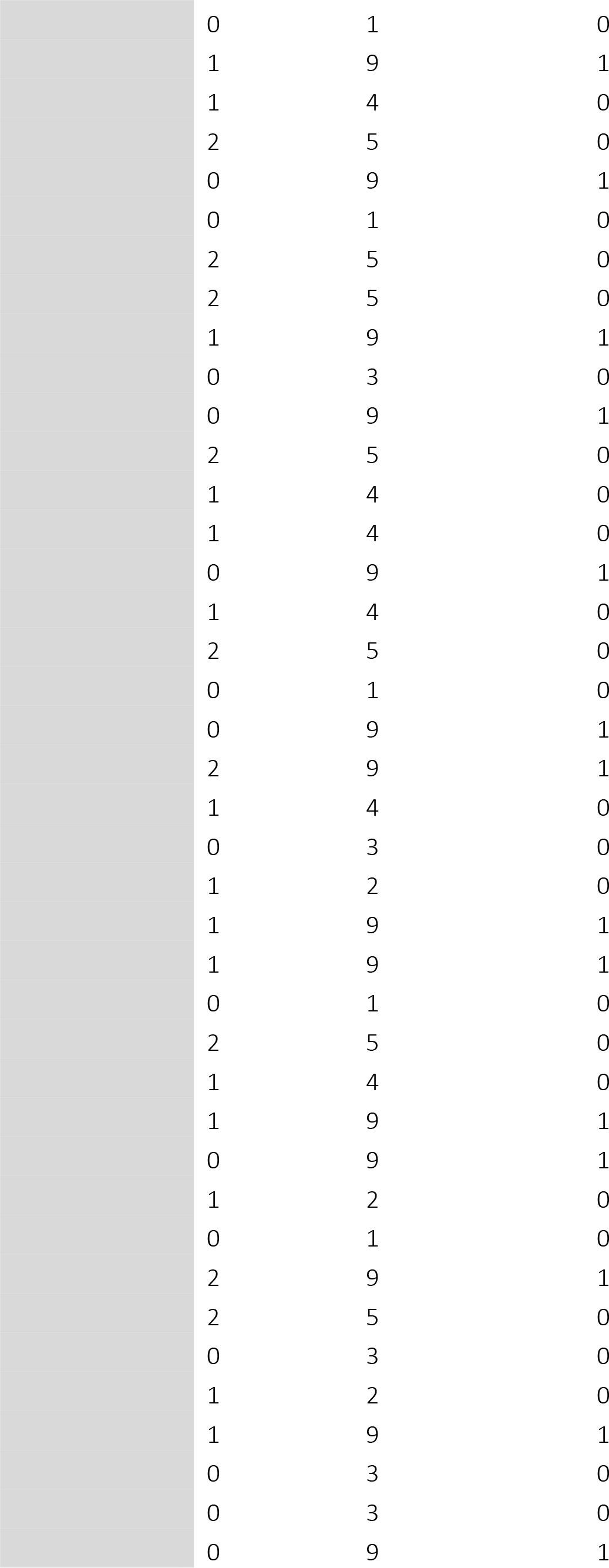
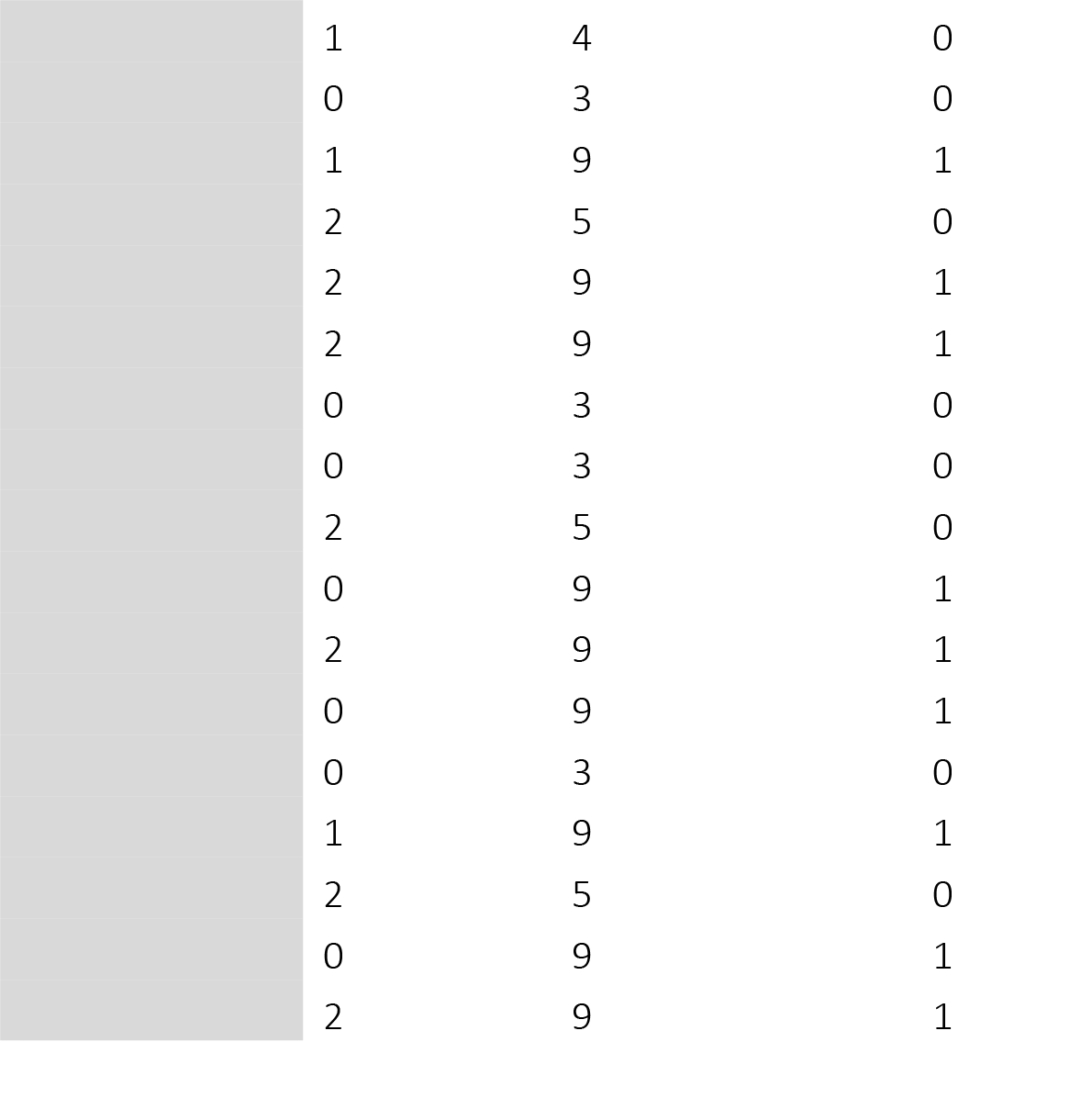
List of word-face pairs used in both Exp. 1 and Exp. 2 categorized according to experimental conditions, together with the presentation order during each run; word emo – emotion category of a word (0 – disgust, 1 – fear, 2 – neutral), face emo – emotion category of a face (0 – disgust, 1 – fear, 2 – neutral), exp cond – experimental condition (1 – disgust congruent, 2 – disgust incongruent, 3 – fear incongruent, 4 – fear congruent, 5 – neutral, 9 – new); novelty – novelty of a word presented during retrieval (0 – old, 1 – new). The list of Polish words is available upon request. Note that the NAWL dataset for Polish was based on the German BAWL- R (Võ et al., 2009), but an English translation of words used in the study can be also available upon request.

**Table S2.** Mean number and proportion of trials with standard deviations for all experimental conditions in Exp. 1 (with ET) and Exp. 2 (with fMRI). M – mean number of trials, SD – standard deviation, % M – mean proportion of trials, % SD – standard deviation of the proportion, dis – disgust, fea – fear, neu – neutral, con – congruent, incon – incongruent, corr – correct, incorr – incorrect.

|  | Exp. 1 (n = 31) |  |  |  | Exp. 2 (n = 37) |  |  |  |
| --- | --- | --- | --- | --- | --- | --- | --- | --- |
|  | M | SD | % M | % SD | M | SD | % M | % SD |
| corr |  |  |  |  |  |  |  |  |
| dis con | 23,26 | 6,41 | 0,58 | 0,16 | 23,86 | 6,32 | 0,60 | 0,16 |
| dis incon | 11,81 | 5,67 | 0,30 | 0,14 | 13,43 | 6,79 | 0,34 | 0,17 |
| fea con | 18,32 | 6,91 | 0,46 | 0,17 | 13,27 | 6,15 | 0,33 | 0,15 |
| fea incon | 13,87 | 5,35 | 0,35 | 0,13 | 19,38 | 6,77 | 0,49 | 0,17 |
| neu con | 45,84 | 14,92 | 0,57 | 0,19 | 49,81 | 11,97 | 0,62 | 0,15 |
| new dis | 46,48 | 12,62 | 0,78 | 0,21 | 43,22 | 12,31 | 0,72 | 0,20 |
| new fea | 46,74 | 12,65 | 0,78 | 0,21 | 41,89 | 12,06 | 0,70 | 0,20 |
| new neu | 44,94 | 12,06 | 0,75 | 0,20 | 41,95 | 11,61 | 0,70 | 0,19 |
| incorr |  |  |  |  |  |  |  |  |
| dis con | 15,55 | 6,28 | 0,39 | 0,16 | 15,08 | 5,88 | 0,38 | 0,15 |
| dis incon | 26,32 | 5,44 | 0,66 | 0,14 | 25,03 | 6,61 | 0,63 | 0,17 |
| fea con | 23,71 | 5,09 | 0,59 | 0,13 | 25,30 | 5,53 | 0,63 | 0,14 |
| fea incon | 19,52 | 6,75 | 0,49 | 0,17 | 19,22 | 6,31 | 0,48 | 0,16 |
| neu con | 31,29 | 14,84 | 0,78 | 0,37 | 28,00 | 11,47 | 0,35 | 0,14 |
| new dis | 11,81 | 12,17 | 0,20 | 0,20 | 15,47 | 11,46 | 0,26 | 0,19 |
| new fea | 12,19 | 11,73 | 0,20 | 0,20 | 16,00 | 11,51 | 0,27 | 0,19 |
| new neu | 13,57 | 11,39 | 0,22 | 0,19 | 16,51 | 11,31 | 0,28 | 0,19 |

**Table S3.**
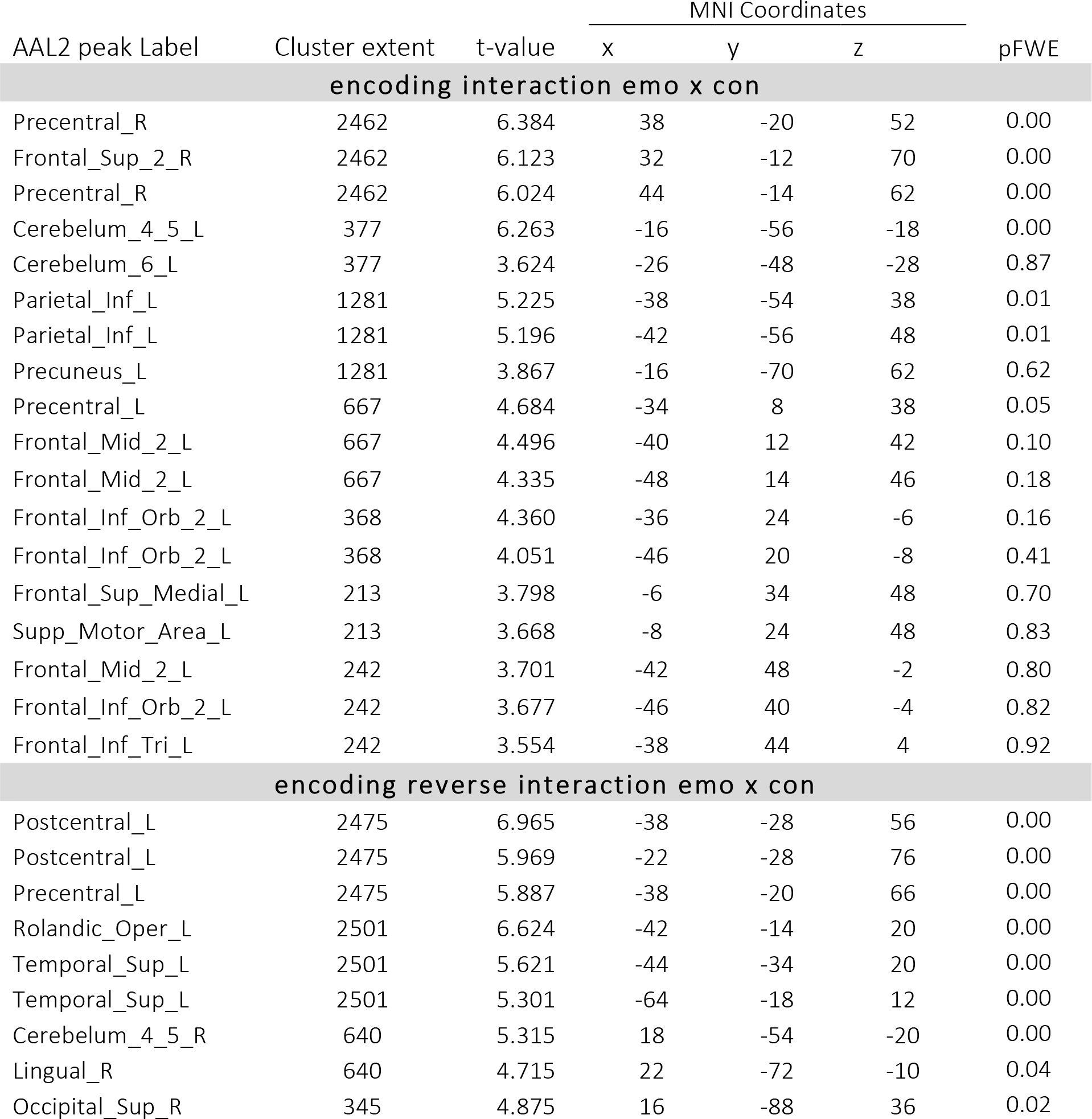
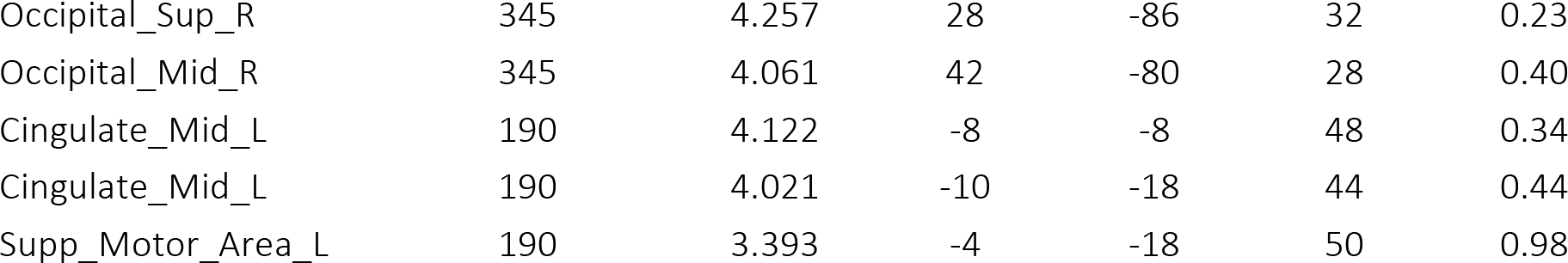
Results of the encoding whole-brain fMRI data univariate analysis (n = 18). A voxel-wise height threshold of p < .001 (uncorrected) combined with cluster-level extent threshold of p < .05 (corrected for multiple comparisons based on Monte Carlo simulation (nn = 2, two-sided, p = .001, alpha = .05, k = 172 voxels) was applied, Table shows all local maxima separated by more than 8mm, regions were automatically labeled using the AAL-2 atlas, x, y, and z Montreal Neurological Institute (MNI) coordinates in the left-right, anterior-posterior, and inferior-superior dimensions, respectively, pFWE signifies p-values at the peak level.

| AAL2 peak Label | Cluster extent | t-value | MNI Coordinates |  |  | pFWE |
| --- | --- | --- | --- | --- | --- | --- |
|  |  |  | x | y | z |  |
| encoding interaction emo x con |  |  |  |  |  |  |
| Precentral_R | 2462 | 6.384 | 38 | -20 | 52 | 0.00 |
| Frontal_Sup_2_R | 2462 | 6.123 | 32 | -12 | 70 | 0.00 |
| Precentral_R | 2462 | 6.024 | 44 | -14 | 62 | 0.00 |
| Cerebelum_4_5_L | 377 | 6.263 | -16 | -56 | -18 | 0.00 |
| Cerebelum_6_L | 377 | 3.624 | -26 | -48 | -28 | 0.87 |
| Parietal_Inf_L | 1281 | 5.225 | -38 | -54 | 38 | 0.01 |
| Parietal_Inf_L | 1281 | 5.196 | -42 | -56 | 48 | 0.01 |
| Precuneus_L | 1281 | 3.867 | -16 | -70 | 62 | 0.62 |
| Precentral_L | 667 | 4.684 | -34 | 8 | 38 | 0.05 |
| Frontal_Mid_2_L | 667 | 4.496 | -40 | 12 | 42 | 0.10 |
| Frontal_Mid_2_L | 667 | 4.335 | -48 | 14 | 46 | 0.18 |
| Frontal_Inf_Orb_2_L | 368 | 4.360 | -36 | 24 | -6 | 0.16 |
| Frontal_Inf_Orb_2_L | 368 | 4.051 | -46 | 20 | -8 | 0.41 |
| Frontal_Sup_Medial_L | 213 | 3.798 | -6 | 34 | 48 | 0.70 |
| Supp_Motor_Area_L | 213 | 3.668 | -8 | 24 | 48 | 0.83 |
| Frontal_Mid_2_L | 242 | 3.701 | -42 | 48 | -2 | 0.80 |
| Frontal_Inf_Orb_2_L | 242 | 3.677 | -46 | 40 | -4 | 0.82 |
| Frontal_Inf_Tri_L | 242 | 3.554 | -38 | 44 | 4 | 0.92 |
| encoding reverse interaction emo x con |  |  |  |  |  |  |
| Postcentral_L | 2475 | 6.965 | -38 | -28 | 56 | 0.00 |
| Postcentral_L | 2475 | 5.969 | -22 | -28 | 76 | 0.00 |
| Precentral_L | 2475 | 5.887 | -38 | -20 | 66 | 0.00 |
| Rolandic_Oper_L | 2501 | 6.624 | -42 | -14 | 20 | 0.00 |
| Temporal_Sup_L | 2501 | 5.621 | -44 | -34 | 20 | 0.00 |
| Temporal_Sup_L | 2501 | 5.301 | -64 | -18 | 12 | 0.00 |
| Cerebelum_4_5_R | 640 | 5.315 | 18 | -54 | -20 | 0.00 |
| Lingual_R | 640 | 4.715 | 22 | -72 | -10 | 0.04 |
| Occipital Sup R | 345 | 4.875 | 16 | -88 | 36 | 0.02 |
| Occipital_Sup_R | 345 | 4.257 | 28 | -86 | 32 | 0.23 |
| Occipital_Mid_R | 345 | 4.061 | 42 | -80 | 28 | 0.40 |
| Cingulate_Mid_L | 190 | 4.122 | -8 | -8 | 48 | 0.34 |
| Cingulate_Mid_L | 190 | 4.021 | -10 | -18 | 44 | 0.44 |
| Supp_Motor_Area_L | 190 | 3.393 | -4 | -18 | 50 | 0.98 |

**Table S4.**
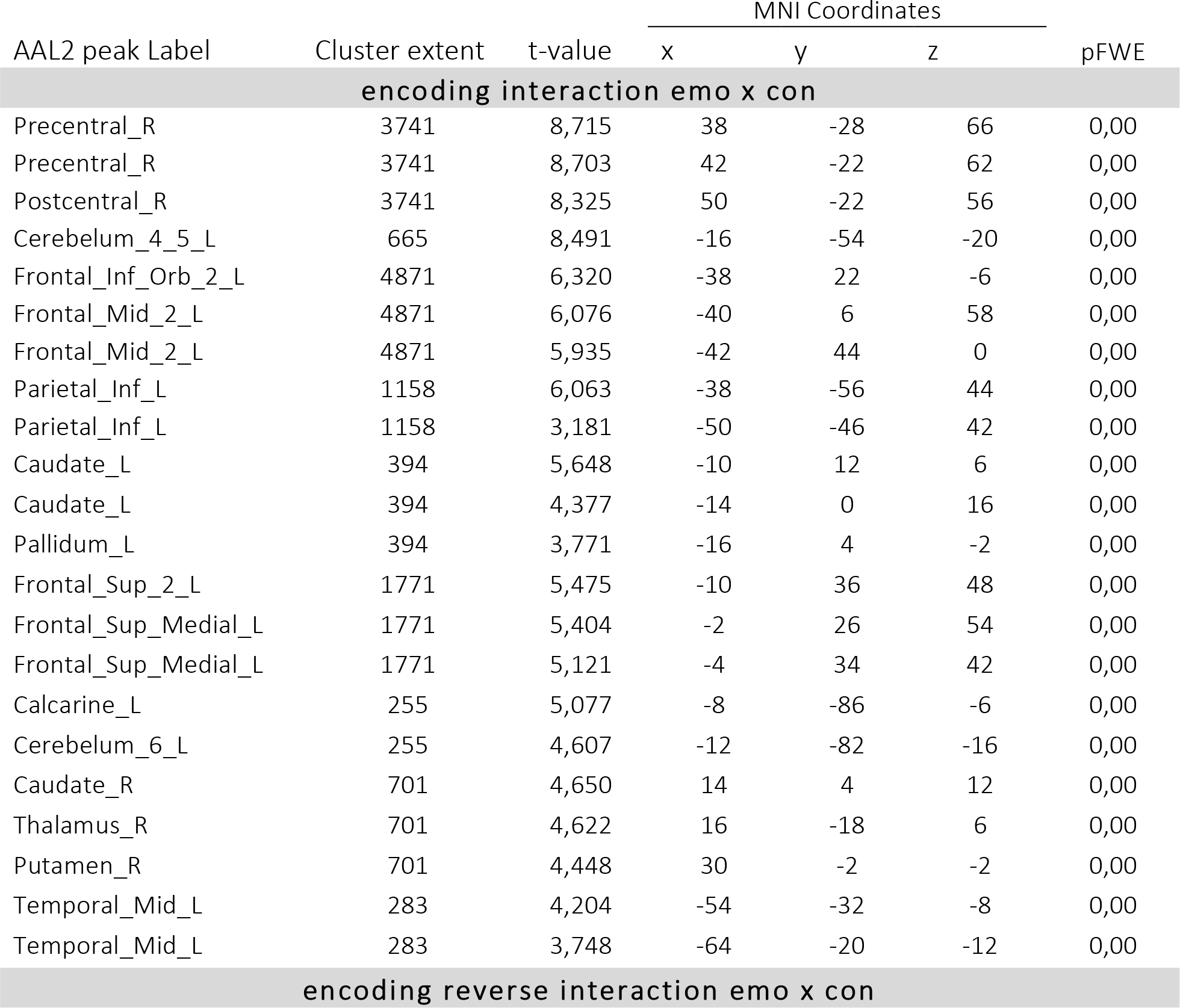
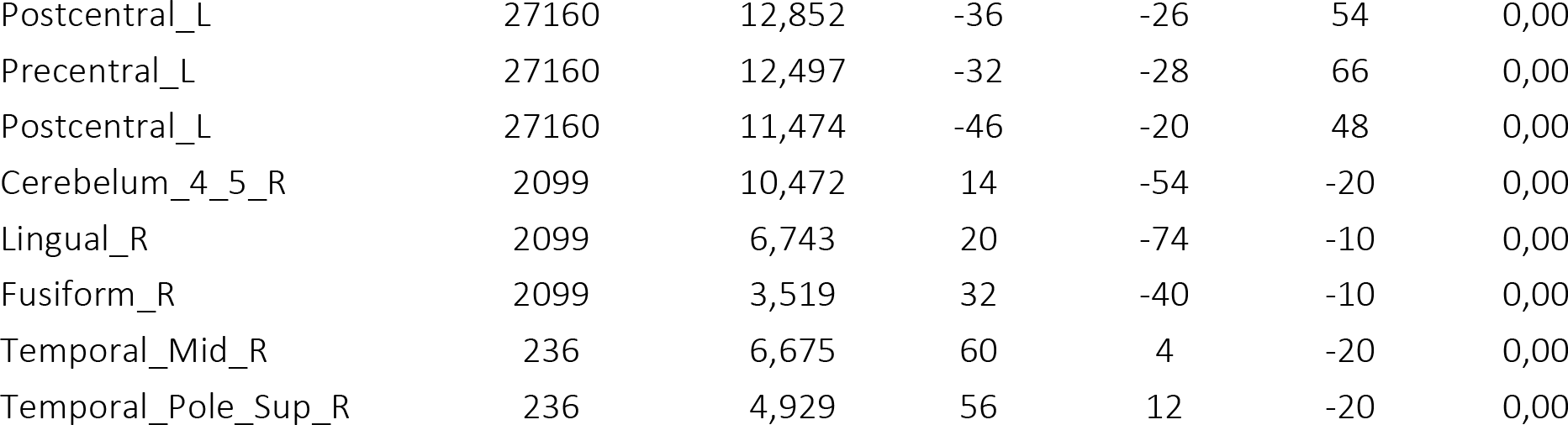
Results of the encoding whole-brain fMRI data univariate analysis (n = 33, only best runs modeled per participant). A voxel-wise height threshold of p < .001 (uncorrected) combined with cluster-level extent threshold of p < .05 (corrected for multiple comparisons based on Monte Carlo simulation (nn = 2, two-sided, p = .001, alpha = .05, k = 172 voxels) was applied, Table shows all local maxima separated by more than 8mm, regions were automatically labeled using the AAL-2 atlas, x, y, and z Montreal Neurological Institute (MNI) coordinates in the left-right, anterior-posterior, and inferior-superior dimensions, respectively, pFWE signifies p-values at the peak level.

**Table S5.** Descriptive statistics of visual characteristics of images (faces) used in the experiment: luminance (an average pixel intensity of the gray-scaled image), contrast (SD of all luminance values) and entropy (a statistical measure of the randomness or noisiness of an image); *N – number of face images per emotion condition, M – mean, SD – standard deviation*

|  |  | N | M | SD |
| --- | --- | --- | --- | --- |
| luminance | disgust | 80 | 103.44 | 26.40 |
|  | fear | 80 | 107.02 | 22.62 |
|  | neutral | 80 | 104.79 | 21.77 |
| contrast | disgust | 80 | 50.26 | 12.44 |
|  | fear | 80 | 49.96 | 12.92 |
|  | neutral | 80 | 49.76 | 11.92 |
| entropy | disgust | 80 | 7.08 | .34 |
|  | fear | 80 | 7.12 | .29 |
|  | neutral | 80 | 7.14 | .28 |

In order to control for the possible effects of visual characteristics of images (faces) used in the experiment, we performed a metadata analysis on the following measures computed for each face image: luminance, contrast and entropy. We run one-way ANOVA to indicate if those measures were equal for the experimental conditions and found that there were no significant differences, neither in luminance [F(2,237) = .466, p = .628], nor contrast [F(2,237) = .033, p = .968], or entropy [F(2,237) = .973, p = .380].

### Memory schemas and emotion schemas

First, based on previous literature, we concluded that ‘memory schemas’ can be defined as “superordinate knowledge structures that reflect abstracted commonalities across multiple experiences, exerting powerful influences over how events are perceived, interpreted, and remembered” (Gilboa & Marlatte, 2017). Thus, one of the critical features of a memory schema is being based on statistical regularities of our experience and much recent literature used term “memory schema” referring to this feature. In most cases, these studies defined memory schemas at the semantic level, but a few studies also investigated more situational memory schemas, based for example on expectancies about typical object locations in virtual reality (Quent, Greve, & Henson, 2021). Given that schema-related terminology has been ambiguous, leading to diverse interpretations, more features have been proposed to distinguish memory schemas from other knowledge structures (Ghosh & Gilboa, 2014). When considering our examples according to this framework, we believe that more defining features of memory schemas apply, sucha as an associative structure (facial expressions and verbal message), being based on multiple episodes (everyday communication), lack of unit detail (no specific face, no specific words) and adaptability (constant updating with each trial).

Second, existing literature led us to a conclusion that “emotion schemas” could be specific examples of memory schemas. Appraisal theories emphasize the situational aspect of emotions, where specific patterns of organism-environment interactions should lead to evolutionary driven, canonical responses (Izard, 2007; Moors, 2020; Russell, 2003). Similarly, emotion schemas are defined as “most frequently occurring emotion experiences, dynamic emotion-cognition interactions that may consist of momentary/ situational responding” that lead to specific behavioural tendencies (Izard, 2009). We believe that communication can be one example of these emotion schemas (Bucci, 2021; Bucci, Maskit, & Murphy, 2016), where co-occurrence of facial and verbal expressions of a particular emotion shapes our reactions and expectations. Ambiguous communication, in turn, would be incongruent with any of these emotion schemas and fail to activate particular reactions.

### Emotion and congruency with schemas modulate memory performance

**Fig. S1.**
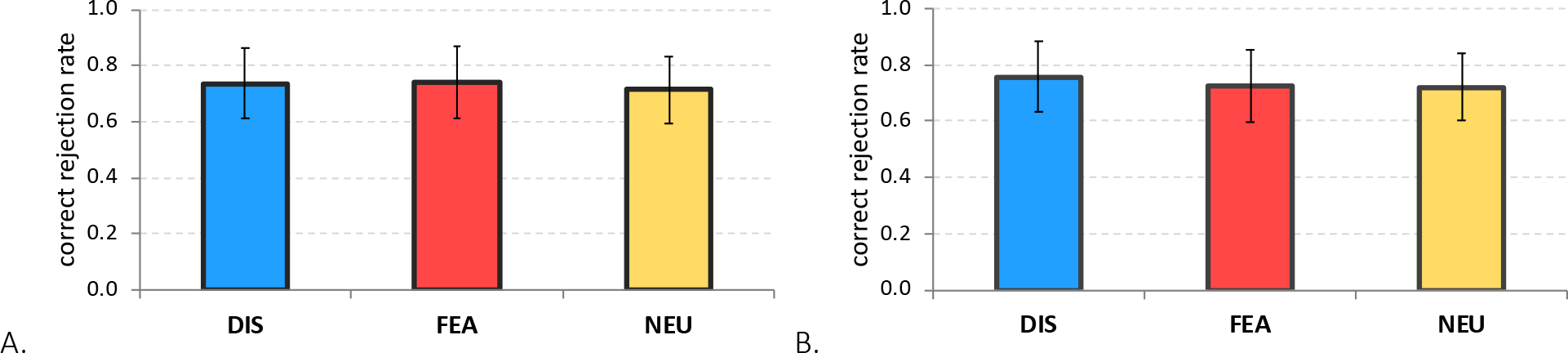
Correct rejection rate (%) for new words across emotions in A. Exp. 1, B. Exp. 2. Colors and labels on horizontal (x) axis indicate retrieved emotion of an associated face: DIS = disgust, FEA = fear; error bars represent one standard deviation.

**Fig. S2.**
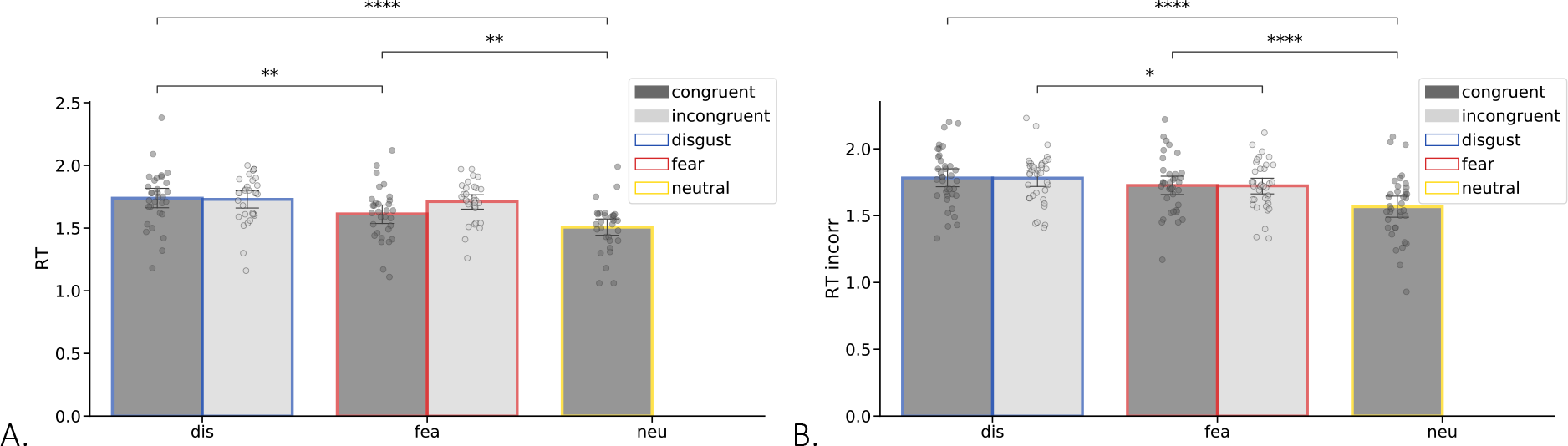
Reaction times (RT) in the retrieval task for *incorrect* retrieval of facial emotions, measured in seconds from the word cue onset until response, across the different emotion and congruency conditions in A. Exp. 1, and B. Exp. 2. *Frame colors and labels on x axis indicate the emotion of associated faces to be retrieved: blue = disgust, red = fear, yellow = neutral; bar colors indicate the congruency with emotion schemas: dark grey – congruent, light grey – incongruent; error bars represent one standard deviation; * p < .05, ** p < .01, *** p < .001, **** p < .0001*.

**Fig. S3.**
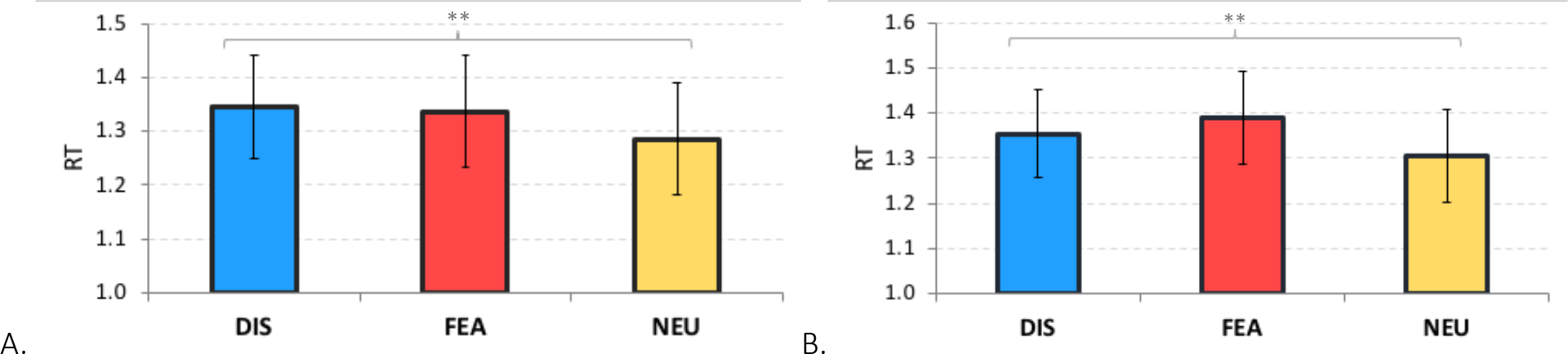
Reaction times (RT) in the retrieval task obtained for correctly rejected new words across emotion conditions in A. Exp. 1, B. Exp. 2. *Colors and labels on horizontal (x) axis indicate retrieved emotion of an associated face: DIS = disgust, FEA = fear; in other words, figure depicts experimental conditions in the following order (according to what was retrieved): congruent disgust, incongruent disgust, incongruent fear, congruent fear; error bars represent one standard deviation; ** p < .01*.

### Reliability of behavioural results across experiments and memory performance

Given that in the fMRI analyses of data from Exp. 2 (with fMRI) we could only include subjects having at least 10 trials per experimental condition (n = 18), we repeated the behavioural analyses for this subset of participants. Similarly, we also analysed a subset subjects having at least 10 trials per experimental condition in Exp. 1 (with ET; n = 16).

In Exp. 1 (with ET, n = 16), memory of facial emotions in response to word cues was tested immediately after encoding. Correct facial emotion memory differed depending on both emotion expression of the face (disgust, fear) and congruency with the word (congruent, incongruent), with a significant interaction effect of these two factors [F(1,30) = 42.963, p < .001, η2 = .741]. Faces from pairs congruent with emotion schemas were generally better retrieved (p < .001). Among congruent pairs, disgust faces were retrieved more often than fear (p = .002), but there was no difference due to emotion among incongruent pairs (p = .146). This result pattern could also mean that disgust words (present in: congruent disgust and incongruent fear conditions) served as a more effective retrieval cue than fear words, possibly enhancing associations with both a disgusted (congruent; p = .002), but not fearful (incongruent; p = .146) face expression. In this case, there was no interaction effect [F(1,30) = 1.608, p = .224, η2 = .097]. In general, disgust words were better cues than fear words [F(1,30) = 42.963, p < .001, η2 = .741] and the cues were better if congruent than incongruent [F(1,30) = 58.547, p < .001, η2 = .790].

In comparison, the neutral faces (always emotionally congruent pairs) were retrieved better than congruent fear faces (p = .003) but equally to congruent disgust faces (p = .252), with a significant effect of emotion across the three old congruent conditions [F(2,30) = 13.556, p < (Widmann, Schröger, Wetzel, et al., 2018).001, η2 = .475]. Correct rejection of new words (i.e. lures during retrieval) did not differ between emotion categories of word cues [F(2,30) = 1.473, p = .246, η2 = .089].

In total, the results of behavioural analyses on a subset of data only from participants having at least 10 trials per condition were largely consistent with the results from the whole samples. This result supports the reliability of behavioural findings and helps the interpretation of fMRI data analysis.

**Fig. S4.**
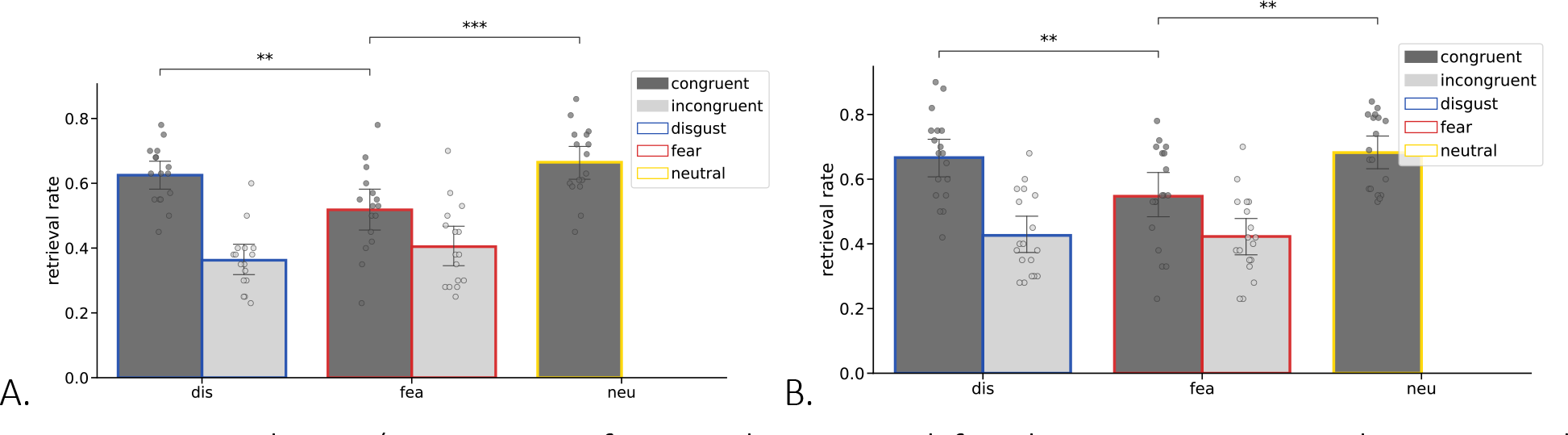
Retrieval rate (proportion of correctly retrieved facial emotions previously associated with old words) across emotion and congruency conditions in A. Exp. 1, and B. Exp. 2, only for participants with >= 10 trials per experimental condition (n = 16 and n = 18, respectively). *Frame colors and labels on x axis indicate the emotion of associated faces to be retrieved: blue = disgust, red = fear, yellow = neutral; bar colors indicate the congruency with emotion schemas: dark grey – congruent, light grey – incongruent; error bars represent one standard deviation, * p < .05, *** p < .001, **** p < .0001*.

### Visual attention to congruent pairs depends on emotion

When comparing how much time during encoding of congruent pairs participants spent looking at ROI1 (face), we found that they looked significantly shorter at disgust faces than fear faces (p < .001) and marginally shorter than neutral faces (p = .054), with a significant effect of emotion on gaze time in ROI1 across the three old congruent conditions [F(2,56) = 11.232, p < .001, η2 = .286]. Emotion modulated also gaze time in ROI2 (word) across the three old congruent conditions [F(2,56) = 3.488, p < .039, η2 = .111], as participants spent marginally more time looking at neutral words than fear-related words (p = .059).

Similarly, emotion of facial expressions in congruent pairs modulated the number of fixations in both ROI1 (face; main effect of emotion [F(2,56) = 7.318, p = .002, η2 = .207]) and ROI2 (word; main effect of emotion [F(2,56) = 7.297, p = .002, η2 = .207]). Similar to the previous analysis, we found more fixations on the face when participants encoded congruent fear than congruent disgust pairs (p = .017) and neutral pairs (p = .006). On the other hand, we observed less fixations on words when encoding congruent fear compared to congruent disgust (p = .008) and neutral pairs (p = .019).

Finally, we found that the number of gaze switches between ROIs as a function of emotion of facial expressions in congruent pairs again showed main effect of emotion [F(2,56) = 12.40, p < .001, η2 = .307]. Participants switched their gaze more between faces and words when encoding congruent disgust pairs than congruent fear pairs (p < .001) and neutral pairs (p = .005).

### Facial emotions modulate pupil dynamics during encoding

When analysing the mean baseline-corrected pupil size in time-windows covering the trial length (0-3s) and comparing changes during encoding as a function of emotion (congruent disgust, fear, neutral) and correctness (correct, incorrect), we found neither a significant interaction effect [F(2,54) = .088, p = .904, η2 = .003], nor the main effect of correctness [F(1,27) = 2.831, p = .104, η2 = .095], but we found the main effect of emotion category [F(1,27) = 5.359, p = .008, η2 = .366]. The pupil change was significantly higher during encoding of congruent disgust pairs than congruent fear pairs (p = .015), but did not differ from neutral pairs (p = .239).

When comparing the pupil PCA component loadings during encoding as a function of emotion (congruent disgust, fear, neutral) and correctness (correct, incorrect), we found neither a significant interaction effect for the late progressive component (1) [F(2,58) = 1.139, p = .322, η2 = .038], nor for the large intermediate transient component (2) [F(2,58) = .454, p = .584, η2 = .015]. However, only for component (1) we found a significant main effect of emotion [F(2,58) = 5.586, p = .009, η2 = .161], and marginally a trend for correctness [F(1,29) = 4.137, p = .051, η^2^ = .125]. The large intermediate transient component (2) loaded higher during encoding of congruent disgust pairs than neutral pairs (p = .002).

### No effect of emotions and emotion schemas in selected brain ROIs during encoding

Reporting results of rm ANOVA on the contrast estimates extracted from each ROI, with emotion of facial expressions (2 levels: disgust, fear) and congruency with emotion schemas (2 levels: congruent, incongruent) as within-subject factors, interaction effects:

a) left HC – n.s. [F(1,17) = .032, p = .86, η2 = .000]

b) right HC – n.s. [F(1,17) = .087, p = .771, η2 = .001]

c) left AMY – n.s. [F(1,17) = .102, p = .753, η2 = .001]

d) right AMY – n.s. [F(1,17) = .006, p = .941, η2 = .000]

Reporting results of rm ANOVA on the contrast estimates extracted from each ROI, with emotion of facial expressions (2 levels: disgust, fear), congruency with emotion schemas (2 levels: congruent, incongruent) and correctness (2 levels: correct, incorrect) as within-subject factors, interaction effects:

a) left HC – n.s. [F(1,17) = .061, p = .808, η2 = .004]

b) right HC – n.s. [F(1,17) = .880, p = .361, η2 = .049]

c) left AMY – n.s. [F(1,17) = .861, p = .366, η2 = .048]

d) right AMY – n.s. [F(1,17) = .076, p = .786, η2 = .004]

Reporting results of rm ANOVA on the contrast estimates extracted from each ROI, with emotion of facial expressions (3 levels: disgust, fear, neutral) as a within-subject factor

a) left HC – n.s. [F(2,34) = .032, p = .86, η2 = .000]

b) right HC – n.s. [F(2,34) = .087, p = .771, η2 = .001]

c) left AMY – n.s. [F(2,34) = .102, p = .753, η2 = .001]

d) right AMY – n.s. [F(2,34) = .006, p = .941, η2 = .000]

**Fig. S5.**
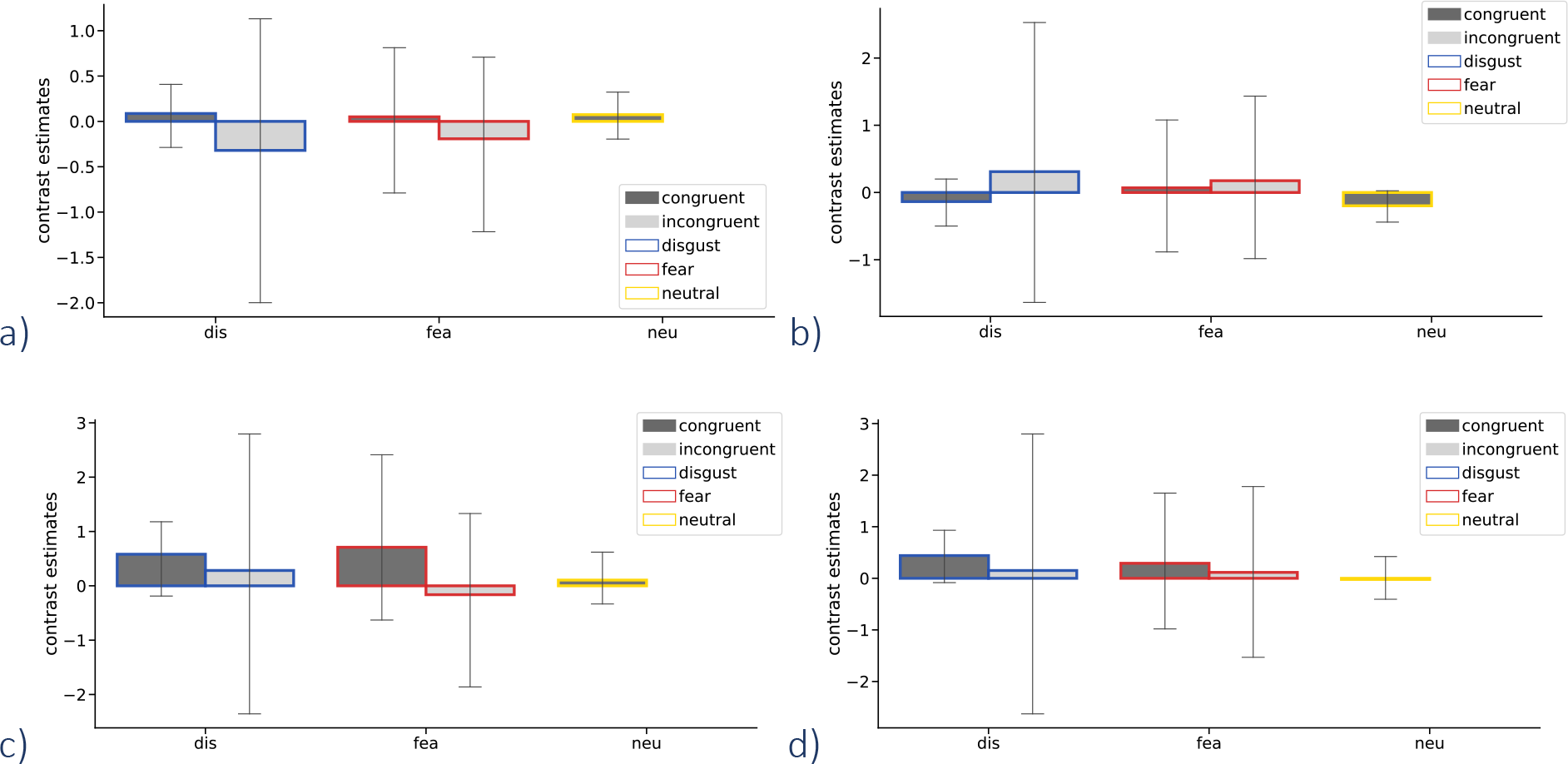
Emotion and congruency effects on the brain activity during encoding. Bars represent contrast estimates for successful encoding (followed by correct retrieval) activity in selected ROIs not reported in the main results (non-significant results): a) left HC, b) right HC, c) left AMY, d) right AMY, split according to emotion and congruency factors (neu condition presented as a point of reference). *Frame colors and labels on x axis indicate the emotion of associated faces to be retrieved: blue = disgust, red = fear, yellow = neutral; bar colors indicate the congruency with emotion schemas: dark grey – congruent, light grey – incongruent; error bars represent one standard deviation*.

An exploratory analysis was also performed to investigate the role of mPFC in emotion and congruency effects. This region was specified according to the AAL2 (Rolls et al., 2015a) atlas as Frontal_Med_Orb and Frontal_Sup_Med (Rolls, Joliot, & Tzourio-Mazoyer, 2015b).

Reporting results of rm ANOVA on the contrast estimates extracted from each explored brain region, with emotion of facial expressions (2 levels: disgust, fear) and congruency with emotion schemas (2 levels: congruent, incongruent) as within-subject factors, interaction effects:

a) left Frontal_Med_Orb - n.s. [F(1,17) = .492, p = .492, η2 = .007) b) right Frontal_Med_Orb - n.s. [F(1,17) = 1.797, p = .198, η2 = .018] c) left Frontal_Sup_Med - trend of significance [F(1,17) = 4.154, p = .057, η2 = .06 d) right Frontal_Sup_Med - n.s. [F(1,17) = .027, p = .87, η2 = .000]

Reporting results of rm ANOVA on the contrast estimates extracted from each explored brain region, with emotion of facial expressions (2 levels: disgust, fear), congruency with emotion schemas (2 levels: congruent, incongruent) and correctness (2 levels: correct, incorrect) as within-subject factors, interaction effects:

a) left Frontal_Med_Orb - n.s. [F(1,17) =.753, p = .398, η2 = .042) b) right Frontal_Med_Orb - n.s. [F(1,17) = 2.788, p = .113, η2 = 141]

c) left Frontal_Sup_Med - trend of significance [F(1,17) = 1.158, p = .297, η2 = .064 d) right Frontal_Sup_Med - n.s. [F(1,17) = 1.055, p = .319, η2 = .058]

Reporting results of rm ANOVA on the contrast estimates extracted from each explored region with emotion of facial expressions (3 levels: disgust, fear, neutral) as a within-subject factor:

a) left Frontal_Med_Orb - n.s. [F(2,34) = .776, p = .405, η2 = .027 b) right Frontal_Med_Orb - n.s. [F(2,34) = 1.774, p = .199, η2 = .054

c) left Frontal_Sup_Med - sig. [F(2,34) = 3.891, p = .047 η2 = .121 d) right Frontal_Sup_Med - n.s. [F(2,34) = .147, p = .772, η2 = .004

Given that we found a significant main effect of emotion in the region of left Frontal_Sup_Med, we performed follow-up post-hoc tests. We found that during successful encoding, there was a greater activation for disgust than fear [t(17) = 2.247, p = .038], and marginally greater than neutral [t(17) = 1.962, p = .066]. However, these results did not survive a correction for multiple comparisons.

**Fig. S6.**
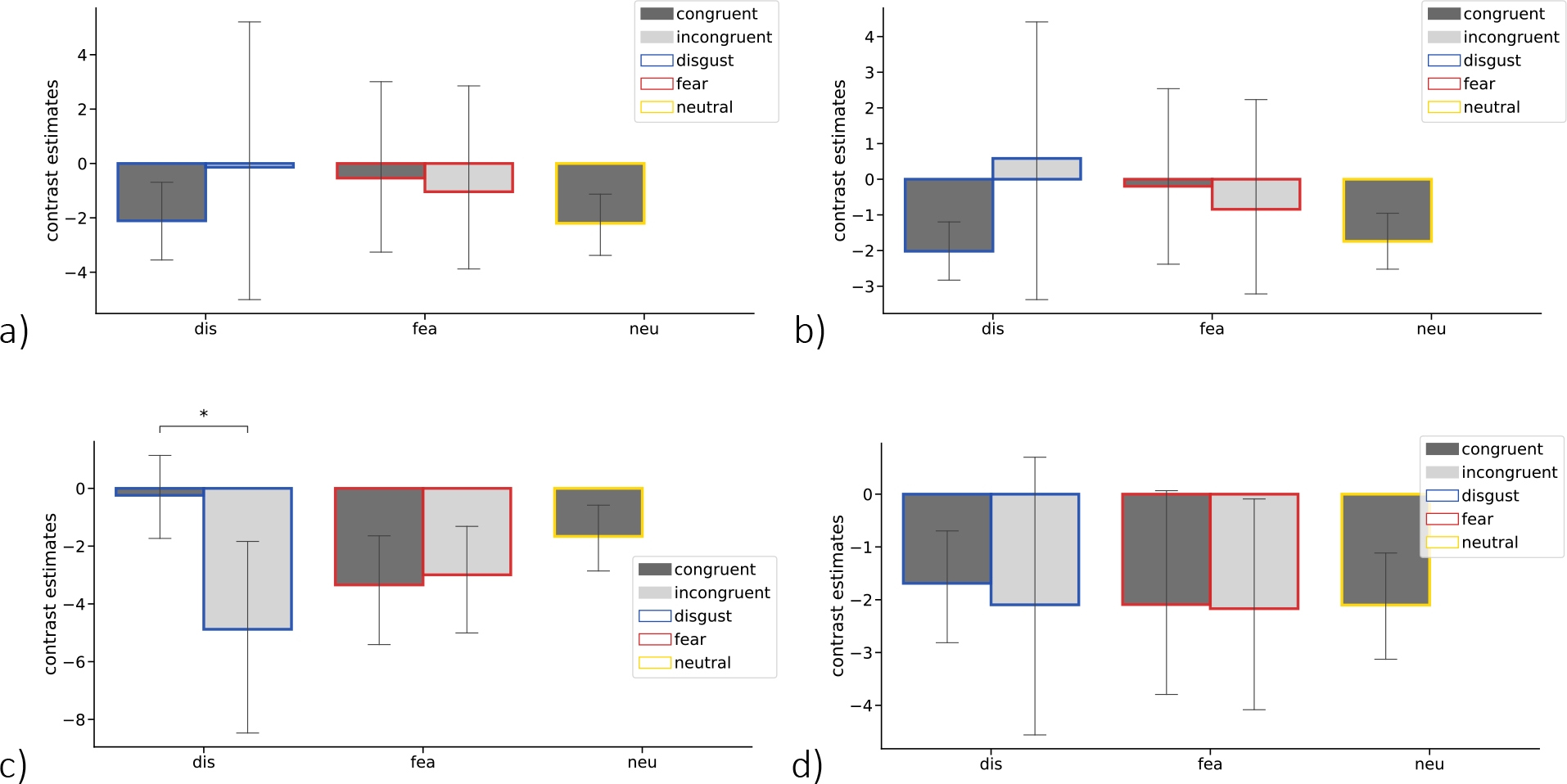
Emotion and congruency effects on the brain activity during encoding. Bars represent contrast estimates for successful encoding (followed by correct retrieval) activity in mPFC – an exploratory analysis not reported in the main results: a) left Frontal_Med_Orb, b) right Frontal_Med_Orb, c) left Frontal_Sup_Med, d) right Frontal_Sup_Med, split according to emotion and congruency factors (neu condition presented as a point of reference). *Frame colors and labels on x axis indicate the emotion of associated faces to be retrieved: blue = disgust, red = fear, yellow = neutral; bar colors indicate the congruency with emotion schemas: dark grey – congruent, light grey – incongruent; error bars represent one standard deviation, * p < .05*.

**Fig. S7.**
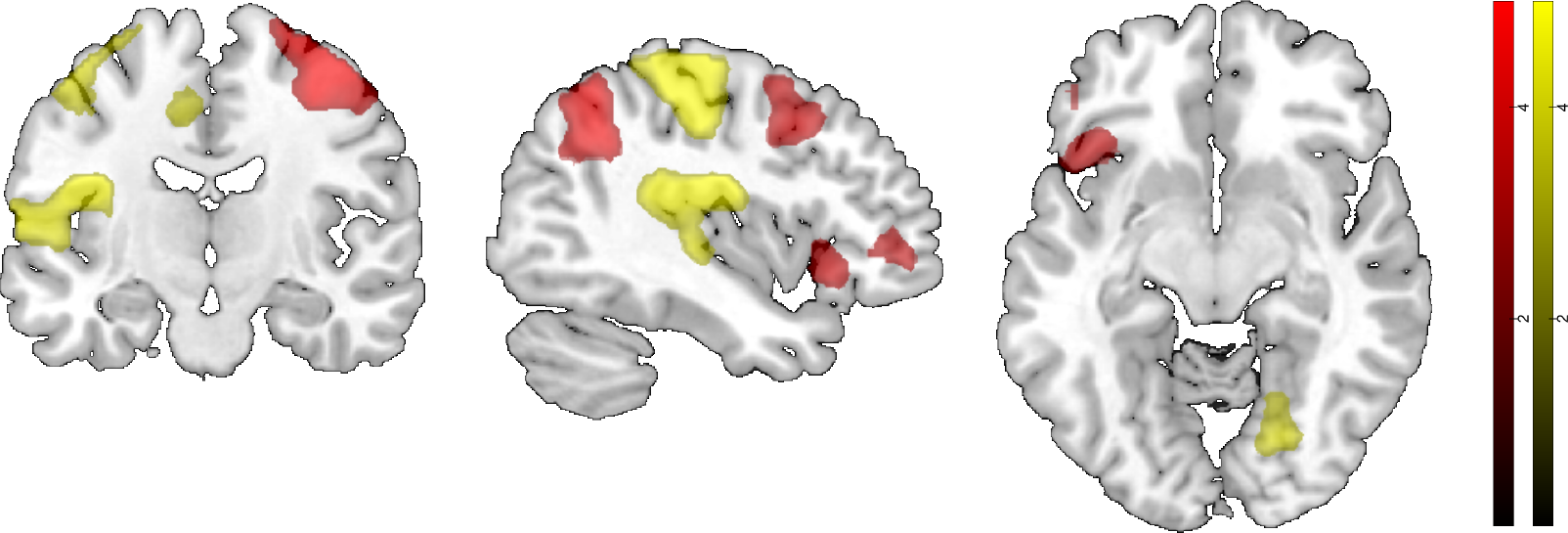
Brain activations showing interaction effects between emotion and congruency during successful encoding (exclusively masked by the same interaction during unsuccessful encoding) in two possible directions: activations higher for disgust > fear and congruent > incongruent (red); activations higher for the reverse contrasts (yellow). Both contrasts contribute to the interaction effect. *Colour scale bars represent a range of t-values; voxel-wise height threshold of p < 0.001 (unc.) combined with a Monte Carlo simulation cluster-level extent threshold (nn = 2, two-sided, p = .001, alpha = .05) of k = 172 voxels. This figure complements the glass brain, Fig. 7a and 7b*.

We made attempts to increase the sample size and performed another individual and group whole brain analysis, this time modeling only the best runs (>= 10 trials per condition) from 33 subjects. Again we tested a positive interaction effect between emotion and congruency during successful encoding, masked by exclusion of the same interaction contrast for unsuccessful encoding. Both analyses (n = 18 and n = 33) yielded similar results, as presented in Table S3 and Table S4. In the “best runs” analysis with an increased sample size, we also observed a higher BOLD signal in left IFG and medial prefrontal cortex, parietal and occipital regions (Fig. S8).

**Fig. S8.**
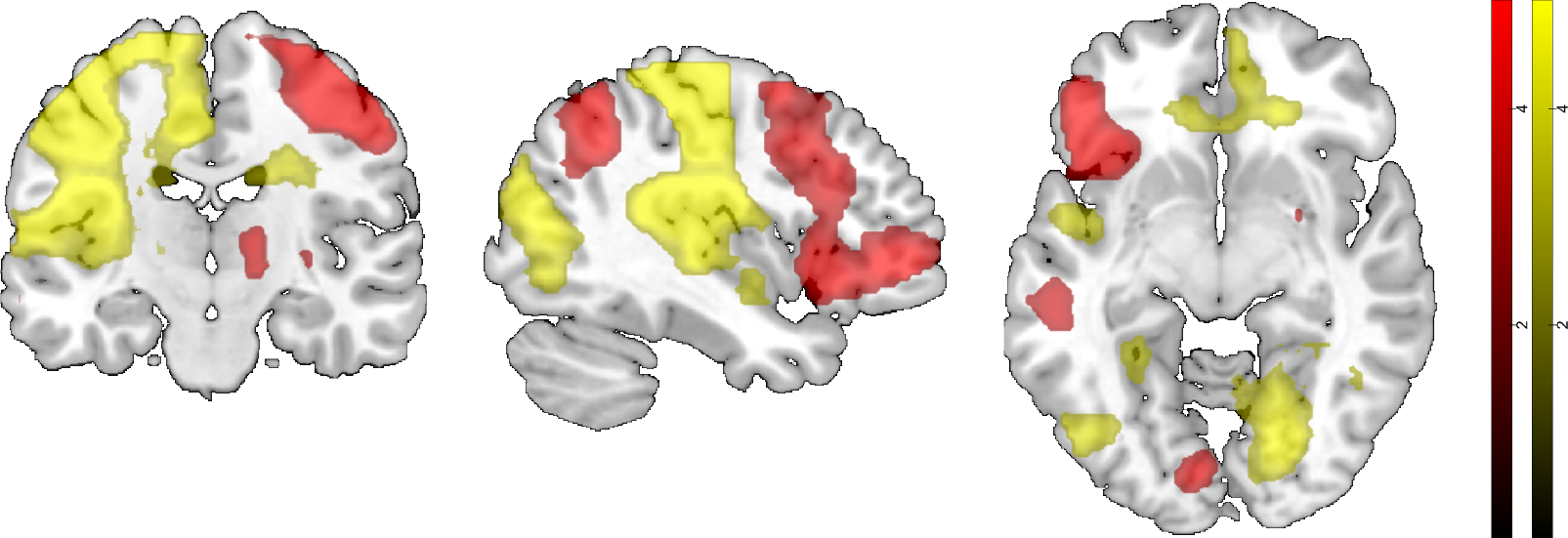
Extended group-level analysis (n = 33, only best runs modeled per participant). Brain activations showing interaction effects between emotion and congruency during successful encoding (exclusively masked by the same interaction during unsuccessful encoding) in two possible directions: activations higher for disgust > fear and congruent > incongruent (red); activations higher for the reverse contrasts (yellow). Both contrasts contribute to the interaction effect. *Colour scale bars represent a range of t-values; voxel-wise height threshold of p < 0.001 (unc.) combined with a Monte Carlo simulation cluster-level extent threshold (nn = 2, two-sided, p = .001, alpha = .05) of k = 172 voxels*.

